# A stronger transcription regulatory circuit of HIV-1C drives the rapid establishment of latency with implications for the direct involvement of Tat

**DOI:** 10.1101/2020.02.20.958892

**Authors:** Sutanuka Chakraborty, Manisha Kabi, Udaykumar Ranga

## Abstract

The magnitude of transcription factor binding site variation emerging in HIV-1C, especially the addition of NF-κB motifs by sequence duplication, makes the examination of transcriptional silence challenging. How can HIV-1 establish and maintain latency despite having a strong LTR? We constructed panels of sub-genomic reporter viral vectors with varying copy numbers of NF-κB motifs (0 to 4 copies) and examined the profile of latency establishment in Jurkat cells. We found surprisingly that the stronger the viral promoter, the faster the latency establishment. Importantly, at the time of commitment to latency and subsequent points, Tat levels in the cell were not limiting. Using highly sensitive strategies, we demonstrate the presence of Tat in the latent cell, recruited to the latent LTR. Our data allude, for the first time, to Tat establishing a negative feedback loop during the late phases of viral infection, leading to the rapid silencing of the viral promoter.

**Importance:** Over the past 10-15 years, HIV-1C has been evolving rapidly towards gaining stronger transcriptional activity by sequence duplication of major transcription factor binding sites. The duplication of NF-κB motifs is unique and exclusive for HIV-1C, a property not shared with any of the other eight HIV-1 genetic families. What mechanism(s) does HIV-1C employ to establish and maintain transcriptional silence despite the presence of a strong promoter and a concomitant strong, positive transcriptional feedback is the primary question we attempted to address in the present manuscript. The role Tat plays in latency reversal is well established. Our work with the most common HIV-1 subtype C (HIV-1C) offers crucial leads towards Tat possessing a dual-role in serving both as transcriptional activator and repressor at different phases of the viral infection of the cell. The leads we offer through the present work have significant implications for HIV-1 cure research.

## Introduction

The post-integration HIV-1 latency is characterized by the presence of the transcriptionally silent but replication-competent provirus within the host cells, a challenge for HIV-1 eradication. A significant amount of controversy surrounds HIV-1 latency following the discovery of a latent HIV-1 reservoir in the resting CD4^+ve^ T-cells (1, 2) whether the external host cell parameters or the intrinsic proviral elements are deterministic in modulating HIV-1 latency. While some models consider HIV-1 latency to be an ‘epiphenomenon’ influenced by the activation status of the host cell and host factors (3–6), others lay emphasis on the stochastic nature of viral gene-expression noise regulated by the LTR-Tat positive feedback circuit, although the influence of the host environmental factors is not disregarded (7–11). Furthermore, several groups have attempted to develop theoretical models to explain the Tat-feedback mediated latency decision in HIV-1 (7, 12).

Master transcriptional regulatory circuits (MTRC) have been identified in prokaryotic and eukaryotic viruses functioning as integral molecular switches to toggle between transcriptional ON and OFF phases. In this context, the lysis-lysogeny decision in bacteriophage λ has been widely researched (13). Fate selection in λ phage is based on the preferential expression of two different key viral proteins, CI and Cro, from a bi-directional promoter, in a mutually exclusive fashion, and on the cooperativity of the CI repressor to establish a ‘bistable’ circuit manifesting lysis or lysogeny (14). Transcriptional circuits in several latency-establishing eukaryotic viruses function through rate-versus-level trade-off where rapid up-regulation of a viral protein is essential for efficient viral replication, but the same molecule is cytotoxic at saturating levels. The immediate-early 2 (IE2) transactivator protein of CMV is a typical example of this phenomenon (15–17). The ICP0 and Rta proteins of Epstein Barr virus (EBV) and Herpes Simplex Virus-1 (HSV-1), respectively, exploit the phenomenon of cooperativity to control alternate replication fates (18–22).

Importantly, the transcription regulatory circuit in HIV-1 appears to differ significantly from those of the λ phage or the eukaryotic viruses mentioned above. First, there is no evidence for a repressor molecule or negative feedback loop controlling the HIV-1 circuit while Tat functioning as only an integral component of a positive feedback circuit. Second, the Tat-feedback circuit seems to lack bistability such that Tat transactivates the LTR as a monomer with no self-cooperativity to form multimers (H=1) (23). Tat was proposed to undergo post-translational modifications at specific sites to modulate latency and to account for Tat mono-stability, and the lack of self-cooperativity. It was proposed that enzymatic conversion of acetylated Tat (Tat_A_) to a more stable deacetylated form (Tat_D_) constitutes a feedback-resistor module with a predominant off state (12). Importantly, the models described above are majorly based on mathematical simulations supported by simple reaction parameters with minimal experimental validation. Of note, these studies exclusively modeled the HIV-1 subtype B system, although other genetic families of HIV-1 contain subtype-specific molecular features.

The examination of transcriptional silence is expected to be technically more challenging in HIV-1C as compared to that in the other subtypes of HIV-1, including HIV-1B, for two different reasons. First, HIV-1C contains several subtype-specific variations in nearly all types of transcription factor binding sites (TFBS) present in the viral promoter, including that of NF-κB, Sp1, RBF-2, and other elements. Among these variations, the copy number difference of the NF-κ B binding elements is the most striking one and unique to HIV-1C. While other subtypes of HIV-1 contain a single (HIV-1E) or two (all others including HIV-1B) NF-κB motifs in the viral enhancer, HIV-1C contains three or four of these motifs (Figure 1). Second, the additional NF-κB binding elements present in HIV-1C are genetically diverse (24). Three different kinds of NF-κB motifs, H, C, and F, may be found in the long-terminal repeat of HIV-1C (C-LTR). We demonstrated previously that the progressive acquisition of the additional NF-κB motifs enhances transcriptional strength of the viral promoter in HIV-1C and confers replication superiority over the canonical viral strains in natural infection, and under experimental conditions (25). Given the positive correlation between the transcriptional strength of the viral promoter and the enhanced strength of transcriptional feedback, it is intriguing how viral latency is favoured in the variant viral strains containing a higher number of NF-κB motifs. In this backdrop, the present study is an attempt to examine the influence of variation in the number of NF-κB binding elements in C-LTR. Of note, the focus of the present study is only on the copy-number difference of the NF-κB binding sites, therefore, on the overall strength of transcription, and its influence on viral latency. The present study does not aim at examining the impact of genetic diversity of NF-κB binding motifs on viral latency.

**Figure 1:**
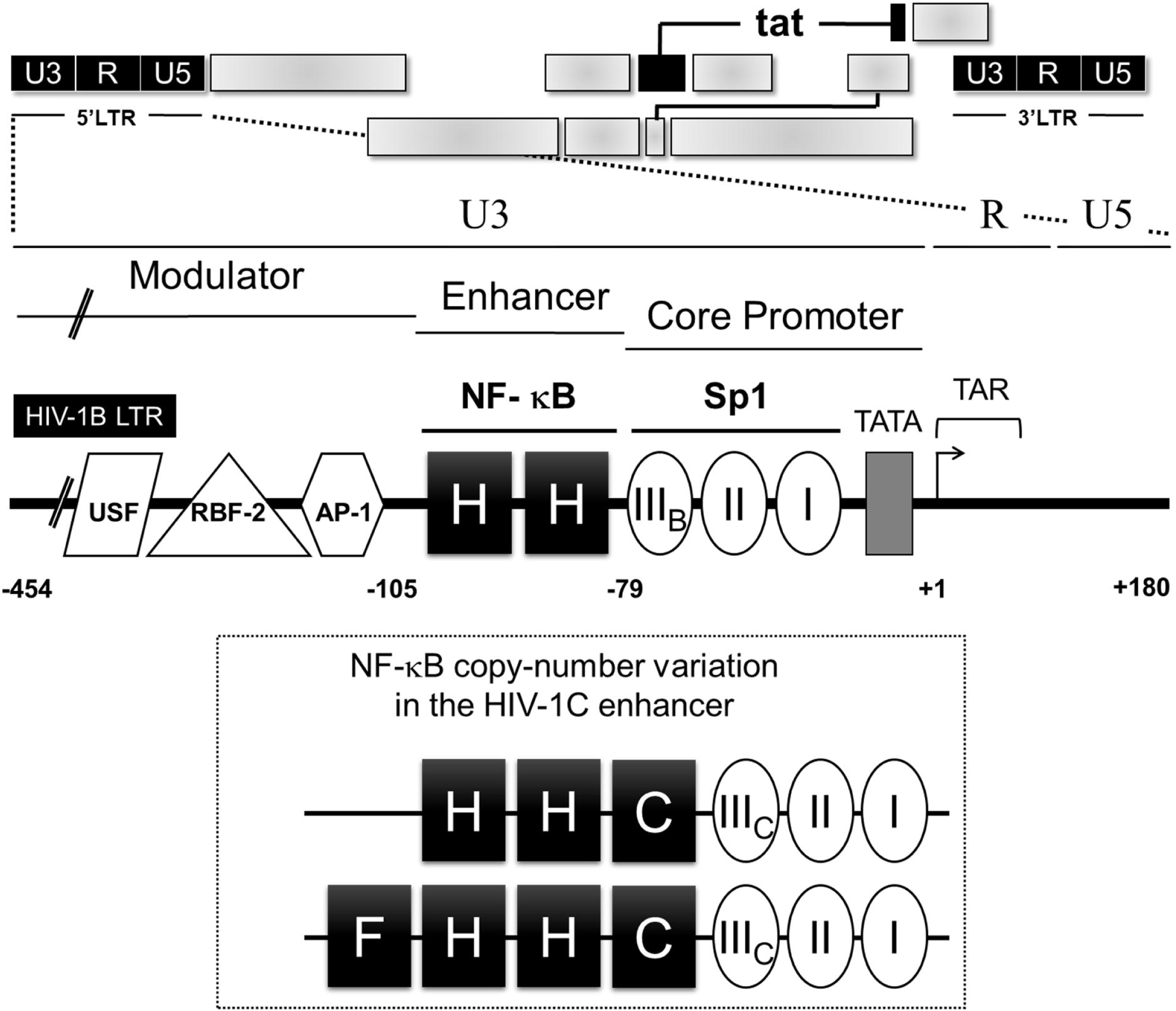
A schematic representation of NF-κB motif diversity in HIV-1C LTR. The canonical HIV-1B LTR containing two identical NF-κB motifs is presented at the top. The distinct regulatory regions (U3, R, U5, the modulator, the enhancer, and the core promoter) and important transcription factor binding sites have been depicted. HIV-1C LTR not only contains more copies of the NF-κB motif (3 or 4 copies) but the additional motifs are also genetically variable (the bottom panel). The three genetically distinct NF-κB motifs present in the C-LTR are denoted as H (GGGACTTTCC), C (GGG<u>G</u>C<u>G</u>TTCC, differences underlined), and F (GGGACTTTC<u>T</u>). Note that the Sp1III site also contains subtype-specific variations as presented; B and C representing respective viral subtypes.

Using sub-genomic HIV-1C reporter viruses that differed in the LTR-Tat transcriptional feedback architecture and using panels of LTR variant viral strains that varied in the copy-number of NF-κB motifs, we demonstrate for the first time that the enhanced transcriptional strength of the LTR leads to a rapid establishment of viral latency. Further, we explain the apparent paradox by demonstrating that a stronger transcriptional activity of the LTR leads to higher levels of cellular Tat protein, which, above a certain threshold, possibly establishes negative feedback on viral transcription. Importantly, using indirect immunofluorescence and a highly sensitive proximity ligation assay, for the first time, we demonstrate the presence of Tat in cells harboring an active or a latent provirus. We also show the recruitment of Tat not only to the active but also the latent proviral LTR, albeit at a magnitude several folds lower. Our data, thus collectively allude to Tat playing a deterministic role in initiating transcriptional silence through negative feedback regulation.

## RESULTS

### The strengths of viral gene expression, as well as Tat-transactivation are directly proportional to the number of functional NF-κB motifs in the HIV-1C enhancer

The present study is an attempt to examine the influence of variation in the number of NF-κB binding elements in the C-LTR on viral gene expression and latency. To this end, we employed two different Jurkat T cell models, the autonomous Tat-feedback (ATF), and the tunable Tat-feedback (TTF) models to examine HIV-1C latency. The sub-genomic viral vectors encoding EGFP (or d2EGFP, a variant GFP with a shorter half-life), were pseudotyped with VSV-G envelope. The two experimental models differ from each other in the manner the LTR-Tat-feedback axis is regulated. Using these two experimental systems, we examined transcriptional activation and silencing as a function of the promoter strength (by varying the copies of NF-κB motifs ranging from 0 to 4 copies in the ATF model) or the Tat-feedback strength (by modulating the physiological concentration of Tat in the TTF model) or both from a panel of HIV-1C LTRs.

The ‘Autonomous Tat-feedback’ (ATF) model of HIV-1 comprises of the presence of only the LTR and Tat, with all the other viral factors being absent, thus retaining the natural functional association between the two major viral factors, as reported previously (7, 26). Several groups have adopted the ATF model to elucidate the mechanisms governing HIV-1 latency. In the present study, we modified the pLGIT sub-genomic reporter vector (7) by substituting the Tat ORF and the 3’ LTR of the parental vector, both of HIV-1B origin, with the homologs of HIV-1C to construct pcLGIT. In the pcLGIT (cLTR-EGFP-IRES-cTat) vector, the expression of EGFP and C-Tat are under the control of the C-LTR.

Given the natural propensity of HIV-1C to contain more copies of the NF-κB motif in the enhancer, three copies typically and up to four copies frequently (24), we constructed a panel of cLGIT viral strains comprising of NF-κB copy-number variant LTRs (p911a series; Materials and Methods). Using the prototype C-LTR containing four functional NF-κB binding sites (FHHC), we introduced inactivating point mutations sequentially into the enhancer to reduce the number of functional NF-κB motifs progressively, from 4 copies to 0 copies (Figure 2A). In some of the subsequent experiments involving the ATF model, we also resorted to the cLdGIT panel of viral strains, where the EGFP reporter was substituted with d2EGFP in the pcLdGIT vector backbone with the same set of NF-κB variant LTRs as the pcLGIT (p911b series; Materials and Methods). The viral stocks of the panel pseudotyped with VSV-G envelope were generated in HEK293T cells, and the relative infectious units (RIU) of the stocks were determined in Jurkat cells using EGFP/d2EGFP fluorescence.

**Figure 2:**
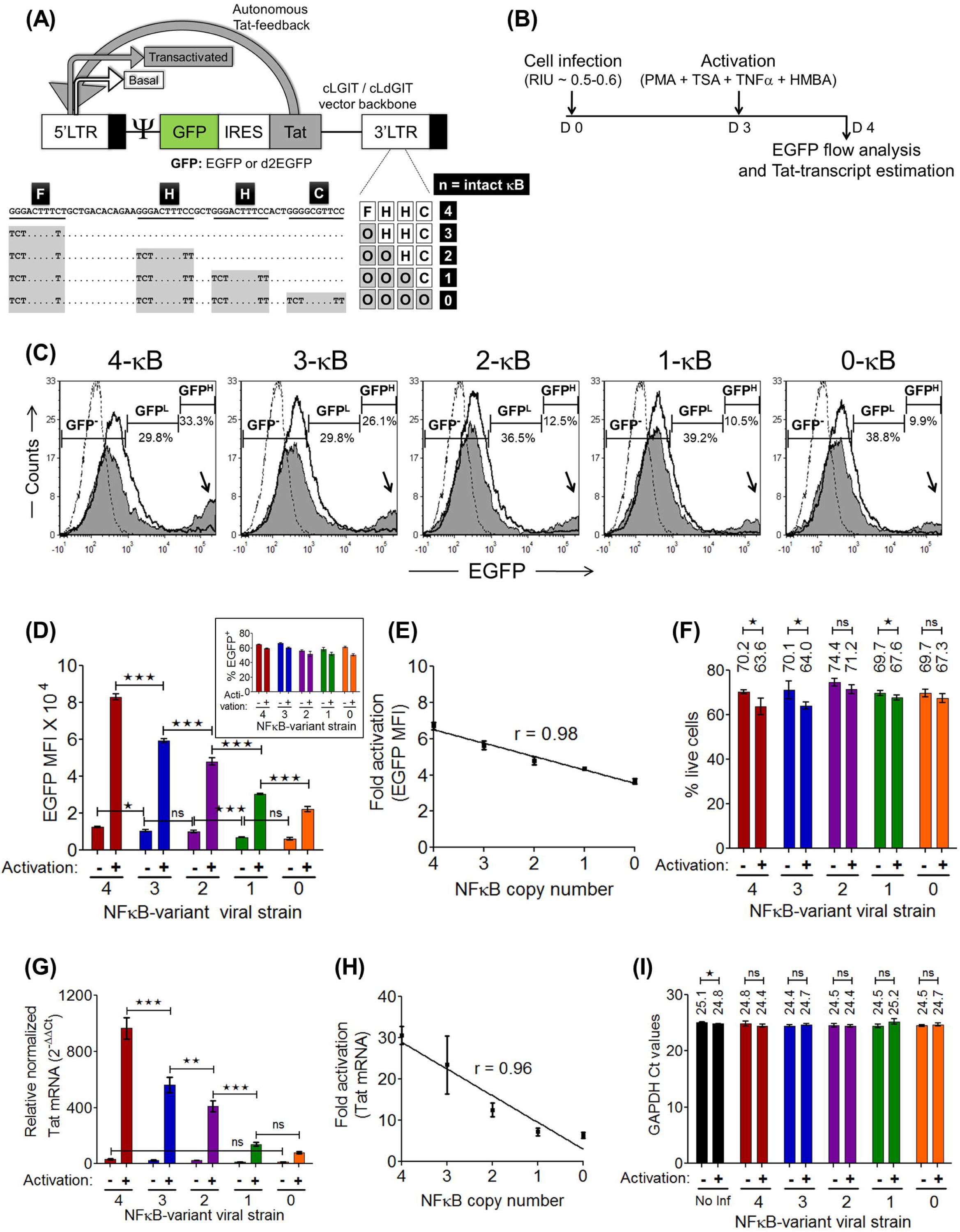
Viral gene expression is proportional to the number of NF-κB motifs in the C-LTR. **(A)** A schematic of the cLGIT/cLdGIT sub-genomic viral vector panels of the autonomous feedback (ATF) model is presented. Note that both the 3’ LTR and Tat are of HIV-1C origin. The nucleotide sequences of the four NF-κB motifs and the inactivating mutations introduced in the motifs of the variant viral strains have been depicted. The EGFP reporter gene (half-life ∼ 48 h) of the cLGIT vector backbone is replaced with its shorter half-life variant d2EGFP (half-life ∼ 2h) in the cLdGIT backbone. **(B)** The time schematic of the gene-expression analysis from the cLGIT vector panel. One million Jurkat cells were infected at an RIU of ∼ 0.5-0.6, independently with each LTR-variant strain. After 72 h of infection, half of the infected cells were activated with a cocktail of global T-cell activators (PMA+TNFα+TSA+HMBA) and 24 h post-activation, EGFP and Tat-transcript expressions were estimated for both the un-activated and activated fractions using a flow-cytometer and a Tat specific RT-PCR, respectively. **(C)** Representative EGFP histograms of the five variant LTRs. The black dotted histogram represents Jurkat cells not infected and not activated; the black hollow histogram represents cells infected but not activated, and the solid grey histogram represents cells infected and activated. The intensity ranges of EGFP^-^, EGFP^Low^, and EGFP^High^ are indicated as GFP^-^, GFP^L^ and GFP^H^, respectively along with the frequency of each fraction (% values). The black arrows point at the gradual decline in the EGFP^High^ (MFI > 10^4^ RFU) population representing the Tat-transactivated population with decreasing copies of NF-κB elements. **(D)** The mean EGFP MFI values from experimental quadruplicates ± SD, data are representative of two independent experiments. Two-way ANOVA with Bonferroni post-test correction was used for the statistical evaluation (*p<0.05, ***p<0.001 and ns – non-significant). The % GFP^+^ profile in the inset confirms near equivalent infection of the target cells with the LTR-variant viral strains at ∼ 0.5-0.6 RIU. **(E)** EGFP-expression manifests a positive linear correlation with the NF-κB copy-number as indicated by the fold enhancement in the EGFP MFI (ratio of the EGFP MFI values of the activated and uninduced fractions from each variant LTR) vs NF-κB copy-number plot. **(F)** Live-dead assay to compare the % live cells between the activated and uninduced pairs. Mean values from experimental quadruplicates ± SD, data are representative of two independent experiments (*p<0.05, ns- non significant; two-tailed, unpaired *t*-test). **(G)** Tat expression was evaluated in an RT-PCR using the ΔΔCt method and GAPDH as the reference gene from total mRNA extracted from 0.2 to 0.5 million cells of un-activated and activated populations. Mean values of the relative Tat-mRNA expression from three independent experiments ± SEM are plotted. Two-way ANOVA with Bonferroni post-test correction was used for statistical analyses. (**p<0.01, ***p<0.001 and ns – non-significant). **(H)** The Tat transcript expression is directly proportional to the NF-κB copy number as observed from the fold enhancement in the Tat transcript levels of the activated fraction over the uninduced fraction. **(I)** Comparable GAPDH Ct values for the different stimulation conditions as well as across the variant viral strains are indicated (*p<0.05, ns- non significant; two-tailed, unpaired *t*-test).

First, we compared the levels of EGFP expression from the LTR-variant cLGIT panel in the context of a functional, positive Tat-feedback loop, where both the reporter gene and the concomitant Tat-feedback strength are expected to vary based on the autoregulatory circuit. Jurkat cells, infected with each viral strain of the cLGIT panel independently at ∼0.5 RIU were either activated with a combination of global T-cell activators (40 ng/ml PMA + 40 ng/ml TNFα + 200 nM TSA + 2.5 mM HMBA) or maintained without activation and 24 hours following the treatment, both the EGFP fluorescence and the Tat transcript levels were examined using flow cytometry and Tat RT-PCR, respectively (Figure 2B). Representative, stacked histograms depicting the three conditions of treatment - uninfected Jurkat cells (black dotted histogram), infected but untreated cells (black hollow histogram) and, infected and activated cells (solid grey histogram) corresponding to all the five NF-κB variant strains are presented (Figure 2C). Importantly, when the cell population in each histogram was demarcated into three categories based on the intensity of EGFP expression (EGFP^-^, EGFP^Low^, and EGFP^High^), it was the EGFP^High^ fraction that displayed the most pronounced impact of the NF-κB site copy number difference on transactivation. The percentage of the EGFP^High^ fraction was directly proportional to the number of NF-κB motifs in the LTR, which was also reflected in the peak height of the EGFP^High^ cluster in the stacked histogram profile.

We quantitated EGFP fluorescence in terms of mean fluorescence intensity (MFI) as a function of the copy numbers of NF-κB motifs in the LTR and found a direct proportionality between them (Figure 2D), although the percent of viral infectivity was comparable (inset; Figure 2D). The LTR containing four NF-κB motifs (FHHC; 4-κB) demonstrated the highest fluorescence intensity with (82,917.51 ± 825.7 RFU) and without (12,365.13 ± 179.3 RFU) activation; while, the LTR in which all the four NF-κB motifs have been mutated (OOOO; 0-κB) demonstrated the lowest levels of the reporter expression with (22,190.38 ± 668.1 RFU) and without (6,083.36 ± 290.5 RFU) activation. The activity of the other three LTRs containing 3 (OHHC; 3-κB), 2 (OOHC; 2-κB), or 1 (OOOC; 1-κB) functional NF-κB motifs remained between the two extremes. The fold enhancement in EGFP expression was directly proportional to the number of functional NF-κB motifs in the LTR with a linear correlation (r = 0.98) between the transcriptional activity and the functional NF-κB motifs in the LTR (Figure 2E). A viability assay performed using a live/dead stain before the analysis of EGFP expression confirmed minimal cell-death following activation (Figure 2F). Similar to the EGFP MFI profile, the level of Tat transcript expression (Figure 2G) and fold transactivation (Figure 2H) were directly proportional to the number of NF-κB copies in the LTR (r = 0.96) with or without activation. The evaluation of the transcripts of GAPDH, as an internal cellular reference control in a real-time RT-PCR, validated the expression levels of Tat mRNA form the viral panel under diverse cell activation conditions (Figure 2I). It is evident from the expression profile that a perfect correlation exists between the number of NF-κB motifs and the level of gene expression from the promoter. Importantly, the expression of EGFP can be used as a surrogate marker for the expression of Tat, since a perfect correlation exists between the two genes co-expressed from the viral promoter. In the subsequent assays, we routinely used the expression of EGFP as a measure of the transcriptional activity of the viral promoter with frequent confirmation of Tat expression.

### A stronger viral promoter establishes latency at a faster rate

A major paradox in the transcriptional regulation of HIV-1C is that a virus that must establish latency tends to acquire a stronger promoter containing more NF-κB motifs, especially when other genetic families of HIV-1 do not employ such a strategy. To understand this paradox, we used the NF-κB copy number variant strains of the ATF panel to determine the kinetics of latency establishment. Using the experimental strategy depicted (Figure 3A), we infected Jurkat cells at a low RIU (∼0.1-0.2) to ensure a single integration event per cell. The cells were allowed to expand before inducing them with a cocktail of global activators, and the EGFP^+^ cells were recovered by sorting. The kinetics of EGFP switch-off was subsequently monitored every four days for 16 days by flow cytometry. A representative strategy of cell gating and sorting is presented (Figure 3B).

**Figure 3:**
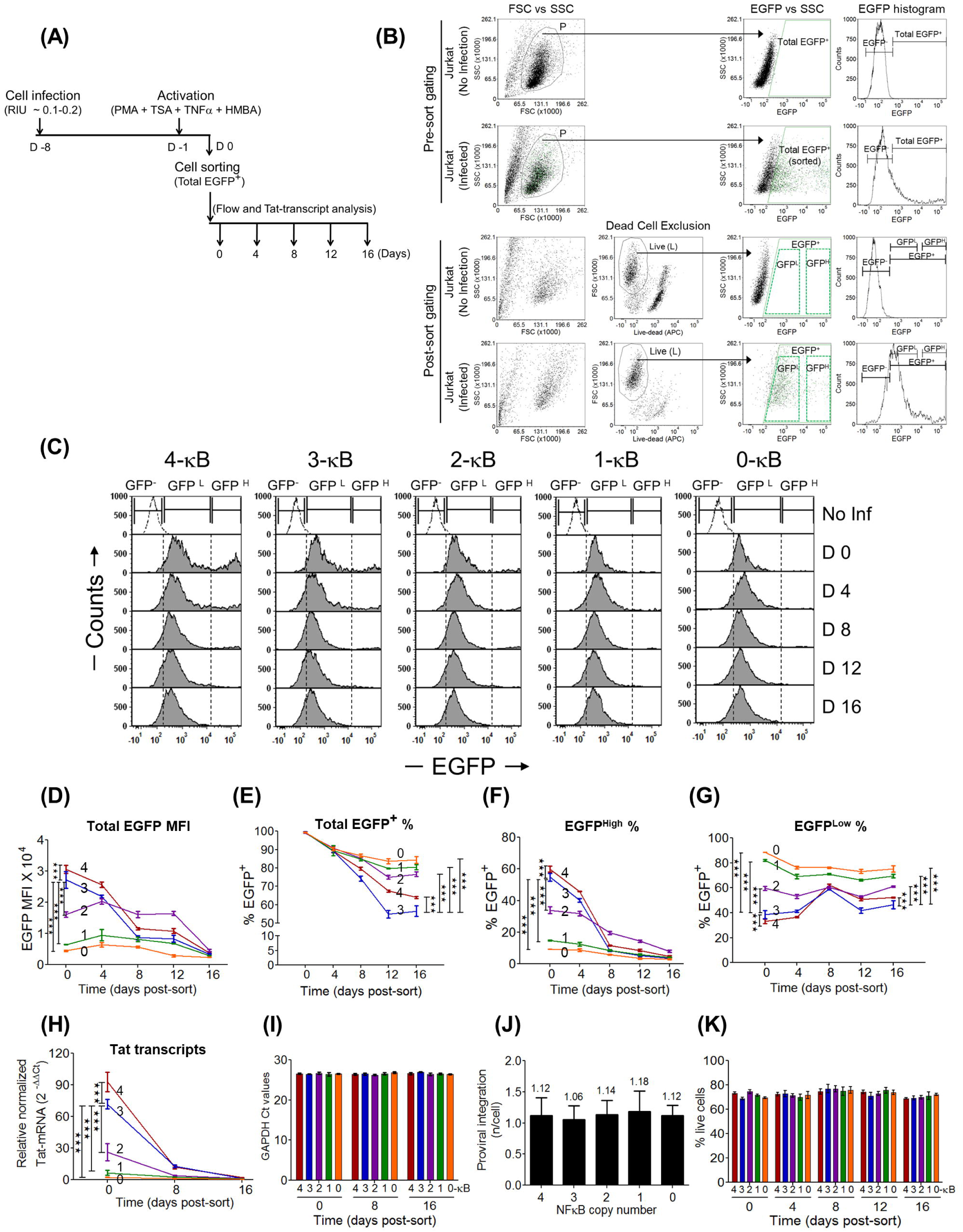
In the ATF model, the stronger the promoter, the faster the latency establishment. **(A)** A schematic representation of the experimental protocol is depicted. One million Jurkat cells were infected with individual strains of the cLGIT panel at a low infectious titer of ∼0.1-0.2 RIU, allowed to relax for a week, treated with the cocktail of global T-cell activators, and 24 h later, all the EGFP^+^ cells (harboring active provirus; MFI >10^3^ RFU) were sorted. The sorted, total EGFP^+^ cells were then maintained in culture, and the EGFP expression was monitored by flow cytometry every four days and that of Tat transcripts on days 0, 8 and 16. **(B)** The gating strategies used to sort the total EGFP^+^ population and for the subsequent latency establishment assay are depicted. Initially, the debris were excluded in the FSC vs SSC scatter plot, and the total EGFP^+^ cells from the population- P were sorted. Importantly, a live-dead exclusion dye was used to stain the post-sorted cells to include only the live, EGFP^+^ for the latency kinetics analyses. The total EGFP^+^ gate was further sub-gated into EGFP^Low^ and EGFP^High^ fractions for subsequent analyses (Figure 3F and 3G, respectively). **(C)** Representative, post-sort, stacked histograms representing the temporal events during transcriptional silencing. The NF-κB variant viral strains demonstrate varying proportions of EGFP^High^ and EGFP^Low^ cells in the total EGFP^+^ sort. **(D), (E), (F),** and **(G)** indicate the kinetic curves corresponding to the total EGFP MFI, percentages of total EGFP^+^, EGFP^High^, and EGFP^Low^ cells, respectively. Data are representative of three independent experiments. Mean values from experimental triplicates ± SD, are plotted. Two-way ANOVA with Bonferroni post-test correction was used for the statistical evaluation (***p<0.001). **(H)** Kinetic curves of relative Tat-mRNA levels of the NF-κB variant strains **(I)** The absolute Ct values of the GAPDH transcripts at different time points. Data are representative of two independent experiments. Mean values from experimental triplicates ± SD plotted. Two-way ANOVA with Bonferroni post-test correction was used for the statistical evaluation (***p<0.001). **(J)** The integration frequency for the five LTR-variant viral strains was estimated using the standard curve and the regression analysis. Viral integration was found to be ∼1.0 per cell for all the five variants. Data are representative of two independent experiments. Mean values from experimental triplicates ± SD values are plotted. **(K)** Live-dead analysis confirms comparable levels of % live cells between the NF-κB variants and across different time-points.

Using the experimental strategy described above, and the cLGIT panel of NF-κB copy number variant viral strains containing 4 to 0 copies of the TFBS, we evaluated how the transcriptional strength of HIV-1 LTR would influence the kinetics of latency establishment over 16 days. The analysis found a profound impact of NF-κB motif copy number on the kinetics of HIV-1 latency establishment. Representative, stacked histogram profiles of the LTR-variant strains depicting the temporal expression pattern of the EGFP^+^ sorted cells are presented (Figure 3C). Although latency establishment was evident for all the five LTRs examined, the rapidity of latency establishment unexpectedly was directly proportional to the number of NF-κB motifs in the viral enhancer (Figures 3D and 3E). In other words, the stronger was the transcriptional activity of the LTR, the faster the latency was established. Based on the slope of EGFP downregulation, the LTRs could be classified into two broader groups: The two strong promoters, the 3- and 4-κB LTRs, down-regulated the EGFP expression at a significantly faster rate than the other three not strong promoters 2-, 1- and 0- κB LTRs. In the present manuscript, we classify the LTRs into two groups, ‘strong’ and ‘weak’ based on the difference in the transcriptional strength, a categorization consistent with many other properties we analyzed subsequently, although the 2- κB LTR occasionally occupied an intermediate position (see below). For instance, the EGFP intensity values (RFU) of 4- κB LTR reduced approximately 8-fold from a value of 30,631.64 ± 1,278.3 on D0 to 3,771.06 ± 245.2 on D16, whereas the corresponding values for the weakest 0- κB LTR were the modest and reduced by only two-folds during the same period from 4,455.11 ± 258.9 to 2,371.98 ± 59.3. Of note, although both 3- and 4- κB LTRs demonstrated a rapid EGFP downregulation, the 3- κB LTR established viral latency at a faster rate, and the difference between the two promoters was highly reproducible and significant. It is not clear if this difference may have implications for the relative replication fitness of the two viral strains.

The expression profile of the Tat transcripts determined using an RT-PCR on days 0, 8 and, 16 also correlated directly with the NF- κB copy number in the LTR, as expected (Figure 3H) and resembled that of the EGFP MFI profile of the LTRs. A profound reduction in Tat expression was observed for all the viral promoters between days 0 and 8. The 4- κB LTR showed the highest level of Tat expression, 92.94 ± 5.4 at D0 that dropped to 12.02 ± 0.8 at D8 and subsequently to 2.3 ± 0.01 at D16. The corresponding values for the 3- κB LTR are 71.76 ± 2.5, 12.8 ± 0.73, and 1.15 ± 0.1, respectively. Of note, comparable expression of GAPDH transcripts was observed from the viral panel at all the time points of latency establishment as quantitated using a real-time PCR (Figure 3I). Furthermore, using a Taqman qPCR, we confirmed a single integration event per cell in all the five stable cell pools, thus ruling out the possibility that the difference in the integration frequency influenced the outcome of the analyses (Figure 3J). A live/dead exclusion assay indicated a comparable percentage of live cells among the panel members, and uniform cell viability was maintained temporally throughout latency establishment (Figure 3K). Importantly, the live/dead gating excluded the dead cells before the EGFP analysis (post-sort gating; Figure 3B); thus, precluding the possibility of EGFP auto-fluorescence from dead cells influencing the data analysis. In summary, our data are suggestive that the enhanced strength of HIV-1C LTR due to the increase in the number of NF-κB sites could play a decisive role in regulating viral latency. A positive correlation between the Tat-transcript levels and the rapid rate of EGFP switch-off by the strong viral promoters is strongly indicative of the Tat-mediated positive feedback loop playing a critical role in establishing viral latency.

### The kinetics of latency establishment is predominantly a function of the EGFP^High^ cells displaying a biphasic mode of transcriptional silence

At the baseline of the above assay, all the variant viral strains were represented by nearly 100% EGFP^+^ cells, but with a varying range of EGFP fluorescence intensities (referred to as total EGFP^+^ cells throughout the manuscript). However, a marked difference in the mean intensity of EGFP among the LTR-variants was noted at D0 time point post-sorting (compare Figures 3D and 3E). This apparent paradox could be explained by analyzing only the EGFP^High^ cells but not the total EGFP^+^ population. In the present essay, we, therefore, gated the cells into two additional subpopulations - EGFP^High^ (MFI >10^4^ RFU) and EGFP^Low^ (MFI ∼10^2^-10^4^ RFU) - as depicted in the post-sort gating strategy (Figure 3B), as evident in the histogram profile of each NF-κB variant strain (Figure 3C). Kinetic curves of % EGFP^High^ and EGFP^Low^ cells were then constructed from the above-gated subpopulations. Importantly, the reduction in the total EGFP MFI (Figure 3D) as well as the Tat-transcript levels (Figure 3H) corresponded perfectly only with the % EGFP^High^ cells (Figure 3F), but not with the % EGFP^Low^ cells (Figure 3G). Thus, the EGFP^High^ cells, not the total EGFP^+^ cells, are decisive in regulating viral latency. Additionally, the % EGFP^High^ temporal curves of the strong (3- and 4-κB) versus weak LTRs (0-, 1- and even 2-κB) were profoundly different. Firstly, on day 0, the strong LTRs produced the highest percentage of EGFP^High^ cells as compared to the weak LTRs. Secondly, the latency establishment of the strong LTRs appeared to have manifested in two distinct phases: a rapid reduction of EGFP expression between days 0 and 8 and a slower rate of decrease after D8; the bi-phase latency profile was either absent or not prominent with the weak LTRs. Thirdly, the rapid fall in EGFP expression of the EGFP^High^ pool of the strong LTRs between days 0 and 8 synchronized with a significant rise in the EGFP^Low^ cell pool peaking on D8. These data collectively allude to the critical role the transcriptional strength of HIV-1 LTR plays in latency establishment. In summary, the EGFP^High^ cell pool, not that of the EGFP^Low^ cells, plays a decisive role in the population latency kinetics of the virus.

### LTR-silencing in the GFP^High^ cells implicates Tat feedback

Given the apparent significance of the EGFP^High^ phenotype for HIV-1 latency establishment, we investigated the phenomenon further by sorting only the EGFP^High^ cell pools for all the NF-κB variant strains that represented a population with a comparable level of fluorescence intensity (Figure 4). At Day 0, the EGFP MFI values were uniform among the variant viral strains of the panel and we monitored downregulation of the green fluorescence every four days for 24 days (Figure 4A). A clear distinction between the strong (4- and 3-κB) and the weak (2-, 1-, and 0- κB) LTRs was evident in the EGFP MFI profile (Figure 4B) or when the EGFP^+^ percentage was considered (Figure 4C), although the 2- κB LTR sometimes occupied an intermediary position; the rate of latency establishment was significantly rapid for strong LTRs. The biphasic mode of latency establishment, rather than a gradual and monophasic mode, was evident from the stacked histogram profiles of the sorted EGFP^High^ pool (Figure 4F). We demonstrated above that a progressively increasing NF-κB site number in the LTR steadily enhances the transcriptional strength as well as the physiological concentration of Tat (Figures 2D and 2G). We, therefore, speculate that higher cellular Tat levels, an invariable outcome of the stronger positive transcriptional feedback, are necessary for the rapid silencing of the LTR as manifested by the EGFP^High^ cells of the strong LTRs.

**Figure 4:**
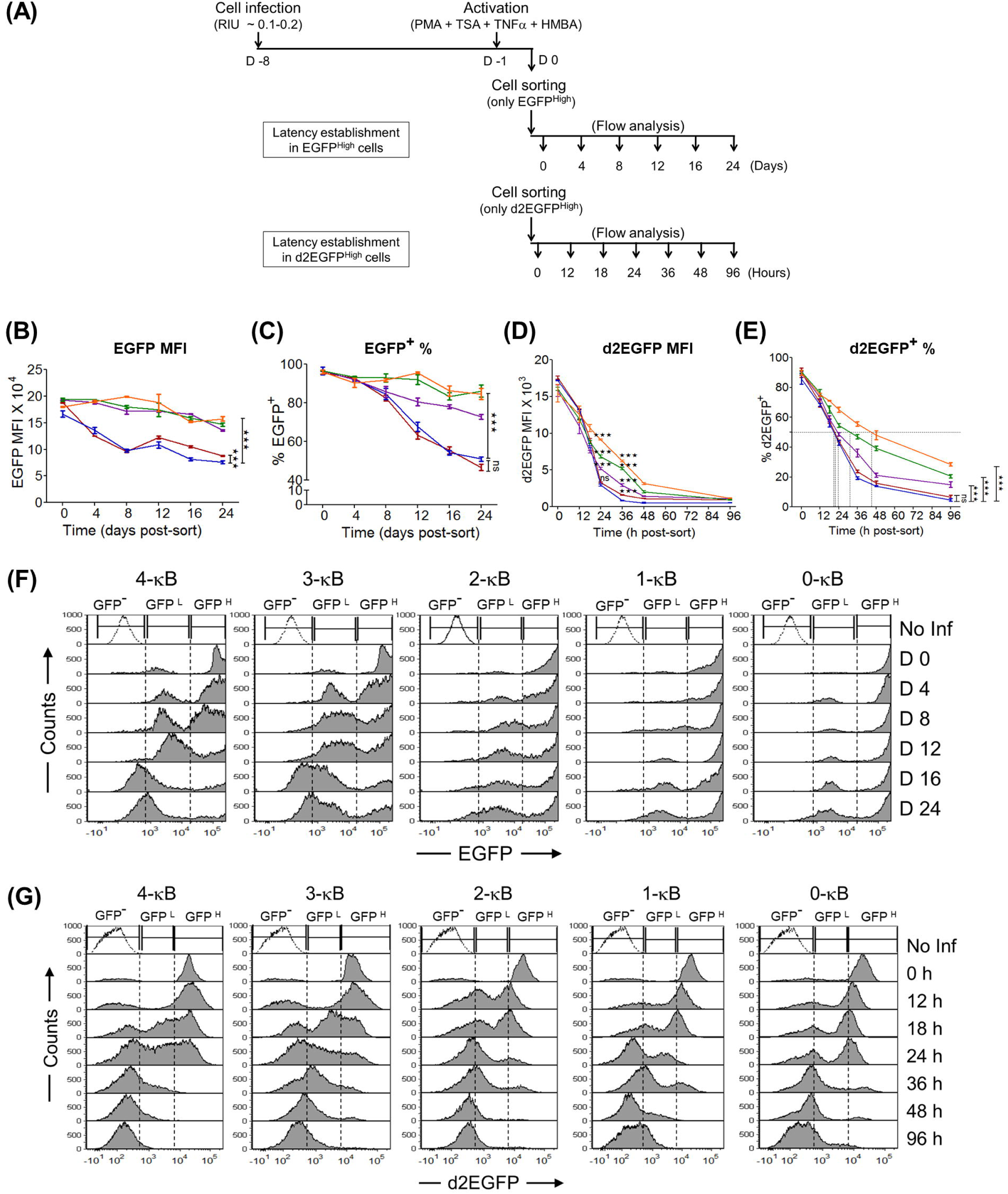
The binary latency trajectory of the GFP^High^ population delineates the viral promoter into strong (3- and 4-κB) and weak (2, 1, and 0-κB) LTRs. **(A)** The experimental schemes to study the kinetics of latency establishment in EGFP^High^ and d2EGFP^High^ cells **(B), (C), (D)** and **(E)** represent the comparative kinetic profiles of EGFP MFI, % EGFP^+^, d2EGFP MFI and % d2EGFP^+^ cells, respectively. Mean values from experimental triplicates ± SD, representative of two independent experiments are plotted. Two-way ANOVA with Bonferroni post-test correction was used for the statistical evaluation (***p<0.001 and ns – non-significant). **(F)** The stacked histogram profiles of the EGFP^High^ cells during latency establishment is presented. The sorted EGFP^High^ cells (GFP^H^; MFI >10^4^ RFU) comprising of a homogeneous population of Tat-mediated transactivated cells, transitioned to the EGFP^-^ phenotype (GFP^-^) through an EGFP^Low^ cluster (GFP^L^; MFI ∼10^2^ – 10^4^ RFU) representing the cells with a basal-level transcription without an intermediate phenotype. **(G)** The stacked histogram profile of the d2EGFP^High^ cells during latency establishment identified regions of d2EGFP^-^ (GFP^-^; MFI <10^3^ RFU), d2EGFP^Low^ (GFP^L^; MFI ∼10^3^–10^4^ RFU), and d2EGFP^High^ (GFP^H^; MFI >10^4^ RFU) phenotypes as demarcated. Of note, given the shorter half-life of d2EGFP, the stability of the d2EGFP^Low^ phenotype was extremely transient; hence the present system lacked a distinct d2EGFP^Low^ cluster at any time point unlike in the EGFP system (Figures 3C and 4G).

Of note, the process of latency establishment above was not complete with any of the LTRs of the cLGIT panel, regardless of the transcriptional strength. The percent of EGFP^+^ cells reached only the halfway mark after 24 days of sorting even for the strong LTRs that established latency at a faster rate (Figures 4B and 4C). Importantly, the long half-life of the EGFP, ∼ 48 h, used in these vectors as a surrogate marker for latency did not represent the actual dynamics of the LTR transcriptional activity faithfully. The cells were continued to be scored as positive for EGFP fluorescence for a significant period even after the LTR was switched off, leading to a false positive scoring. To rectify this problem, we substituted EGFP in the reporter viral strains with d2EGFP characterized by a significantly shorter half-life (2 vs. 48 h) (27). The viral strains of the new panel (cLdGIT) are analogous to the previous panel.

Using the new panel, we sorted the d2EGFP^High^ cells as above to establish the profiles of latency. Several differences in the profiles of latency were readily evident between the cLGIT and cLdGIT panels (compare Figures 4B and 4D; 4C and 4E). Unlike the cLGIT panel, the cLdGIT variants successfully established a near-complete viral latency, and all the members of the panel demonstrated latency establishment at a faster rate; the d2EGFP MFI values reduced to the baseline within 96 h following sorting (Figure 4D). Although the substitution of EGFP with d2EGFP masked the differences in latency kinetics among the members of the cLdGIT panel to some extent, the overall pattern of latency establishment was consistent with that of the cLGIT counterparts. The percentage of the cells downregulating d2EGFP expression was directly proportional to the number of the NF-κB motifs in the viral promoter (Figure 4E). For instance, the time required for the loss of fluorescence in half of the cells (FL_50_) was estimated to be 23.3, 22.1, 24.64, 32.9, and 48 h for the 4-, 3-, 2-, 1-, and 0-κB viral strains, respectively. Thus, a direct correlation between the transcriptional strength of the viral promoter and the rate of latency establishment was consistent between the cLdGIT and cLGIT panels. The bi-phasic mode of latency establishment was also evident in the cLdGIT model (Figure 4G).

Collectively, our data are assertive that the transcriptional strength of the HIV-1 promoter is an essential regulatory parameter for viral latency. Further, the latency kinetics in the Tat-transactivated population (GFP^High^ cells), of two different models, cLGIT and cLdGIT vectors of the ATF panel, followed an NF-κB-site copy number-dependent transcriptional silencing.

### A bimodal (ON or OFF) latency establishment in the pools of cloned cell lines

The observation that the transcriptional strength of the LTR and the feedback loop of Tat function synergistically to silence the viral promoter was drawn based on cell pools. Since individual cells in a pool are heterogeneous in several biological properties, including the site of proviral integration, we examined the nature of the latency profile in multiple cloned cell lines of all the five LTR variants. Jurkat cells were infected with the viral strains of cLGIT (ATF) panel, stimulated with the global activation cocktail, single EGFP^High^ cells were sorted into individual wells of a 96-well culture plate, the sorted cells were allowed to expand for 3-4 weeks, and the EGFP expression profiles were assessed by flow cytometry (Figure 5A). We recovered 16-25 clones from each NF-κB variant, and 16 clones from each variant were randomly selected for the latency analysis. Of note, since each cell line descended from a single parental cell, all the daughter cells derived from the parental cell are expected to have a common site of integration.

**Figure 5:**
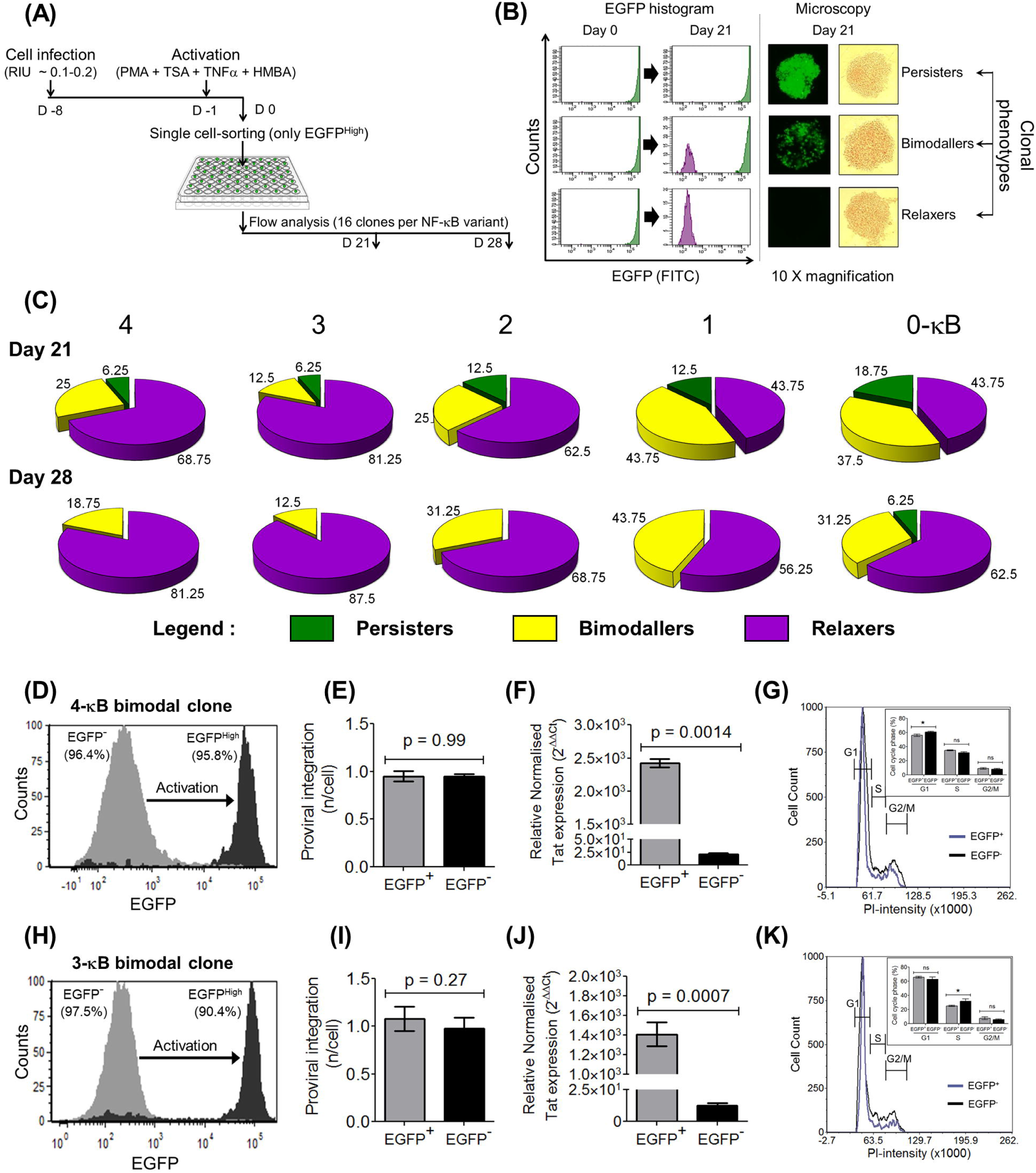
The manifestation of three distinct latency phenotypes of single EGFP^High^ cells. **(A)** The experimental layout of latency establishment in single-cell clones is essentially similar to that of the non-clonal, population kinetics as in Figure 4A. The EGFP^High^ cells (MFI >10^4^ RFU) were single-cell sorted, expanded for three-four weeks, and the pattern of EGFP expression was assessed on days 21 and 28 post-sorting by flow cytometry and fluorescence microscopy. EGFP expression of 16 randomly selected clones, corresponding to each viral variant was measured. **(B)** Based on the fluorescence profile, three distinct categories of clone-persisters (EGFP^High^, MFI >10^4^ RFU), relaxers (EGFP^-^, MFI <10^3^), and bimodallers (binary population of persisters and relaxers) were identified. **(C)** The relative proportion of the above three phenotypes among the LTR-variants as a function of time as indicated at Day 21 and 28 post-sorting. **(D)** and **(H)** Two bimodal cell lines, 3c and 8c representing the 4-κB and 3-κB LTRs, respectively, demonstrate bimodal gene expression. The sorted EGFP^-^ cells from the 4-κB and 3-κB clones generate 95.8% (Figure 5D) and 90.4% (Figure 5H) EGFP^High^ cells, respectively, following activation with the global activation cocktail with negligible proportion of cells displaying the intermediate phenotype. **(E)** and **(I)** A Taqman qPCR targeting a region of the LTR, performed as in Figure 3J, confirms a comparable integration frequency (∼1.0 provirus per cell) between the EGFP^-^ and EGFP^High^ fractions in both the 4-κB (Figure 5E) and the 3-κB (Figure 5I) clones. **(F)** and **(J)** A quantitative real-time PCR for the Tat-transcripts demonstrated significantly higher levels of Tat-transcripts in the EGFP^High^ fraction compared to that in the EGFP^-^ fraction for both the clones. Mean values from three independent experiments ± SEM are plotted. A two-tailed, unpaired *t*-test was used for the statistical evaluation. **(G)** and **(K)** DNA cell-cycle analysis was performed on the EGFP^-^ and EGFP^High^ sub-fractions of the 4-κB (Figure 5G) and the 3-κB (Figure 5K) clones following the standard PI-staining protocol. The overlay histograms for the two subfractions showing the G1, S and G2/M phases are presented. The proportions of cells in the G1, S and G2/M stages were calculated for both the subfractions and depicted in the inset. Mean values from triplicates, representative of two independent reactions ± SD are plotted. A two-tailed, unpaired *t*-test was used for the statistical evaluation.

Based on the EGFP expression pattern, the clones could be categorized into three distinct types (Figure 5B). The persisters, all the daughter cells descending from a single parental cell sustain expression of high-intensity EGFP throughout the observation period of 28 days and even beyond, comparable to that of the original parental cell, indicative of a provirus transcribing actively in all the daughter cells. The relaxers, all the daughter cells of the EGFP^High^ parental cell, have switched-off EGFP expression entirely during the period of observation. The bimodallers, the third clonal type, demonstrated a distinctive feature of the simultaneous existence of both the phenotypes among the daughter cells, although all the cells in the cluster were derived from the same EGFP^High^ parental cell. One subset of the cells maintained high EGFP expression (EGFP^High^), whereas the other subset down-regulated the reporter gene completely (EGFP^-^) with the minimal manifestation of an intermediate phenotype.

Importantly, all the five viral strains of the panel displayed the three clonal phenotypes described above with the distinction that the proportion of the three phenotypes is directly correlated with the copy number of NF-κB sites in the LTR. Given the limitation of available cells for the flow analysis, we could determine the phenotype of the clonal cells only at D21 and D28, not earlier. We analyzed 16 randomly selected clones for each of the five LTRs of the panel (Figure 5C). The profile of the three phenotypes varied significantly among the members of the panel and appeared to associate with the transcriptional strength of the LTR. On D21, a larger proportion of cell lines representing the strong viral promoters (4- and 3-κB LTRs) transited to the OFF state as compared to those of weak promoters (1- and 0-κB LTRs); in contrast, the cells of 2- κB LTR occupied an intermediate position. On D28, the strong LTRs contained no persistent phenotype and fewer bimodal clones. The strong viral promoters downregulated EGFP expression, both persistent and bimodal phenotypes at a significantly faster rate as compared to the other three promoter variants. Despite the limitation of the small number of clonal cell lines used in the analysis, these data are broadly consistent with the results of cell pools (Figure 4). Thus, cell pools and clonal cell populations, both the models demonstrated a direct correlation between the transcriptional strength of the LTR and the rate of latency establishment. Further, both the experimental models are also consistent with each other in demonstrating a bimodal, not a gradual, latency establishment.

The clonal cell lines that display the bimodal EGFP phenotype offer an excellent experimental model as these clonal lines demonstrate two contrasting phenotypes (EGFP^High^ and EGFP^-^ expression) despite an identical viral genotype, chromatin background and, host-cell activation. We selected two clones, 3c and 8c representing the strong 4-κB and 3- κB LTRs, respectively, and characterized them for their bimodality. The two daughter populations, EGFP^High^ and the EGFP^-^ were subsequently enriched using FACS sorting and examined for the nature of transcription complexes recruited to the active and silent LTRs using ChIP analyses (see below). Importantly, a vast majority of the EGFP^-^ cells of the bimodal clones representing the 4- or 3-κB LTRs could be fully reactivated to the EGFP^High^ phenotype; 95.6% (Figure 5D) and 90.4% (Figure 5H) EGFP^High^ cells, respectively, following global activation. The levels of proviral integration between the two subpopulations of each bimodal clone were comparable and close to ∼1.0 (Figures 5E and 5I), ruling out the possibility of integration frequency differences underlying the bimodal phenotype. Importantly, the Tat transcript levels in the EGFP^+^ subfractions of both the clonal cell lines were significantly higher compared to their EGFP^-^ counterparts, approximately 112 folds for the 4-κB (Figure 5F) and 80 folds for the 3-κB clones (Figure 5J). We also excluded the possibility of the bimodal phenotype arising from cell cycle differences between the two phenotypes. We compared the proportion of cells in the different phases of the cell cycle (G1, S, and G2/M) between the EGFP^-^ and EGFP^High^ subpopulations. We found that the cell proportions were comparable in both the phenotypes. The data were reproducible in both the 4- (Figure 5G) and the 3-κB (Figure 5K) clones; the manifestation of the two contrasting phenotypes in the bimodallers was therefore unlikely to be a consequence of cell-cycle differences.

### A tunable regulatory circuit of HIV-1 transcription alludes to the direct role of Tat in latency establishment

The stronger transcriptional activity of the LTR is expected to lead to a proportionately higher expression of Tat, which in turn should increase the transcriptional activity of the LTR further. As a consequence of the unique arrangement, the two principal regulatory elements collectively modulate viral gene expression. In this backdrop, the profile of latency kinetics observed using the ATF model above cannot be ascribed to the different functional activity of either of the elements alone. It was, therefore, necessary to employ a strategy where Tat transactivation alone becomes a variable factor while the transcriptional strength of the LTR remains constant. To this end, we constructed a new HIV-1-Jurkat cell line model, the ‘Tunable Tat-feedback’ (TTF) model, where the transactivation strength of Tat can be modulated independently while keeping the transcriptional strength of the LTR constant. Tat in the TTF model was engineered to possess two unique properties as compared to that in the ATF model (Figure 6A). First, Tat was fused with DsRed2-RFP (stated as RFP throughout the manuscript) to express as a fusion protein enabling the direct visualization of its expression. The new HIV-1 reporter vector pcLdGITRD (cLTR-d2EGFP-IRES-Tat:RFP:DD), thus, co-expressed two different fluorescent proteins, d2EGFP and Tat-RFP, under the control of the LTR. Second, the Tat-RFP fusion protein was tagged with the C-terminal degradation domain (DD) of FK506 binding protein (Tat:RFP:DD). The DD domain can target the fusion protein for rapid proteasome-mediated degradation (28). Shield1, a small molecule ligand, however, can rescue the DD-mediated degradation by specifically interacting with the DD motif and stabilizing the target protein in a dose-responsive manner (29). The sub-genomic HIV-1 reporter vector pcLdGITRD, representing the TTF model, thus, can fine-tune the intracellular concentration of the ‘Tat:RFP:DD’ fusion protein by changing the concentration of Shield1 in the culture medium, in the context of a fixed LTR strength.

**Figure 6:**
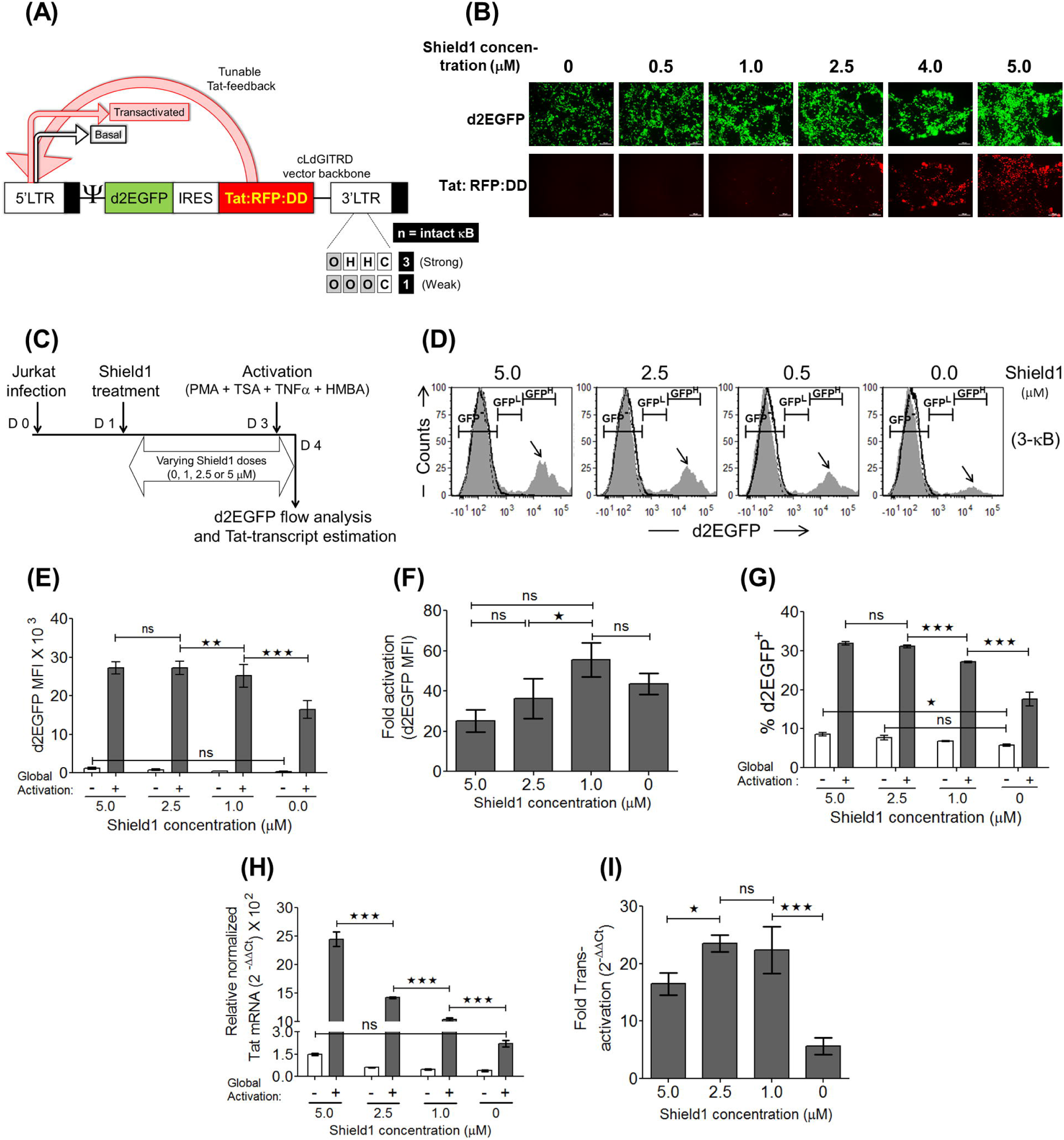
In the tunable Tat-feedback (TTF) model, the stronger the LTR-Tat feedback, the higher the viral gene expression. **(A)** A schematic of the sub-genomic HIV-1 vector backbone- cLdGITRD representing the TTF model. The 3- and 1-κB LTRs representing a strong and a weak promoter, respectively, have been used in the present study. The small molecule Shield1 stabilizes the ‘Tat:RFP:DD’ cassette in a dose-dependent fashion making it available for the subsequent transactivation events at the LTR. **(B)** Validation of Shield1 dose-dependent stabilization of the ‘Tat:RFP:DD’ cassette in HEK293T cells. One mg of the cLdGITRD-3-κB vector was transfected into 0.6 million HEK293T cells in separate wells in the presence of varying concentrations of Shield1 as indicated, and the images were captured 48 h post-transfection. The experiment was repeated twice with comparable results. **(C)** The experimental layout to confirm Shield1 dose-dependent gene expression and Tat-transactivation in Jurkat cells is presented. Approximately 0.3 million Jurkat cells were infected with the LdGITRD-3-κB strain (20 ng/ml p24 equivalent), and post 24 h, the infected cells were split into four fractions, each treated with a different concentration of Shield1 as indicated. After 48 h of Shield1 treatment, half of the cells from each fraction were activated for 24 h followed by the quantitation of d2EGFP and Tat-mRNA expression levels for both the induced and uninduced fractions. **(D)** Representative stacked histograms indicate a Shield1 dose-dependent Tat-transactivated population (black arrows) at a fixed LTR-strength (fixed number of NF-κB motifs). The peak-height of the d2EGFP^High^ population (MFI > 10^4^ MFU) proportionally reduced with the Shield1 dose; the TTF model thus confirmed the d2EGFP^High^ cluster to represent the Tat-transactivated population. **(E)** The Shield1 concentration dependent d2EGFP MFI, **(F)** fold d2EGFP enhancement, and **(G)** % d2EGFP^+^ plots are presented. Values from experimental triplicates ± SD, representing two independent experiments are plotted. Two-way ANOVA with Bonferroni post-test correction was used for the statistical evaluation (**p<0.01, ***p<0.001 and ns – non-significant). **(H)** Relative Tat-transcript levels and **(I)** fold Tat-mediated transactivation were evaluated as in Figures 2G and 2H, respectively. The mean values from three independent experiments ± SEM are plotted. Two-way ANOVA with Bonferroni post-test correction was used for the statistical evaluation (*p<0.05, ***p<0.001 and ns – non-significant).

We constructed a panel (cLdGITRD, the p913 series; Materials and Methods) of two LTR-variant viral strains consisting of 3 or 1 NF-κB motifs, representing the strong and weak LTRs, respectively (Figure 6A). A direct correlation between the Shield1 concentration in the medium, ranging from 0 to 5 μM, and the intensity of Tat-RFP expression was observed in HEK293T cells using the 3-κB viral reporter vector (Figure 6B). Importantly, the viruses could infect the target Jurkat cells. We evaluated the levels of d2EGFP expression and the Tat-mediated transactivation, with an increasing concentration of Shield1 in the medium, using the experimental strategy as depicted (Figure 6C). Interestingly, the effect of Shield1 concentration was directly manifested on the d2EGFP^High^ population in the stacked histogram profile (black arrow), indicating Shield1 dose-dependent Tat transactivation and also confirming that the d2EGFP^High^ phenotype represented the Tat-transactivated cells (Figure 6D). A direct correlation was also established in the stable Jurkat cells between the Shield1 concentration and d2EGFP MFI or ‘Tat:RFP:DD’ expression (Figures 6E and 6H respectively) suggesting Shield1-dependent stabilization of the ‘Tat:RFP:DD’ cassette and the subsequent Tat-mediated LTR transactivation. Of note, although we normalized the viral infection, the % d2EGFP^+^ values demonstrated a dose-response proportional to the Shield1 concentration even though the d2EGFP itself does not contain the DD of FKBP (Figure 6G). The optimal fold activation of the d2EGFP expression (Figure 6F) and Tat transcript levels (Figure 6I) were found to be 1 μM and 2.5 μM, respectively. In the subsequent experiments, therefore, we used Shield1 in the range of 0 to 3 μM.

Importantly, the fusion of Tat with DsRed2-RFP offered the advantage of tracking the expression of Tat in real-time during latency establishment. To determine the kinetics of latency establishment in Jurkat cells, we used an experimental schematic as depicted (Figure 7A). Jurkat cells were infected with 3- or 1-κB viral strain at an RIU of ∼0.1-0.2 in the presence of 1 μM Shield1 and expanded for a week in the presence of Shield1. Subsequently, the cells were activated with the global activators for 24 h, the d2EGFP^High^ population (MFI ∼10^4^ RFU) was sorted, the sorted cells were maintained separately at four different concentrations of Shield1 (0, 0.5, 1.0 and 3.0 μM), and the levels of d2EGFP and Tat-RFP expression were monitored every 24 h by flow cytometry.

**Figure 7:**
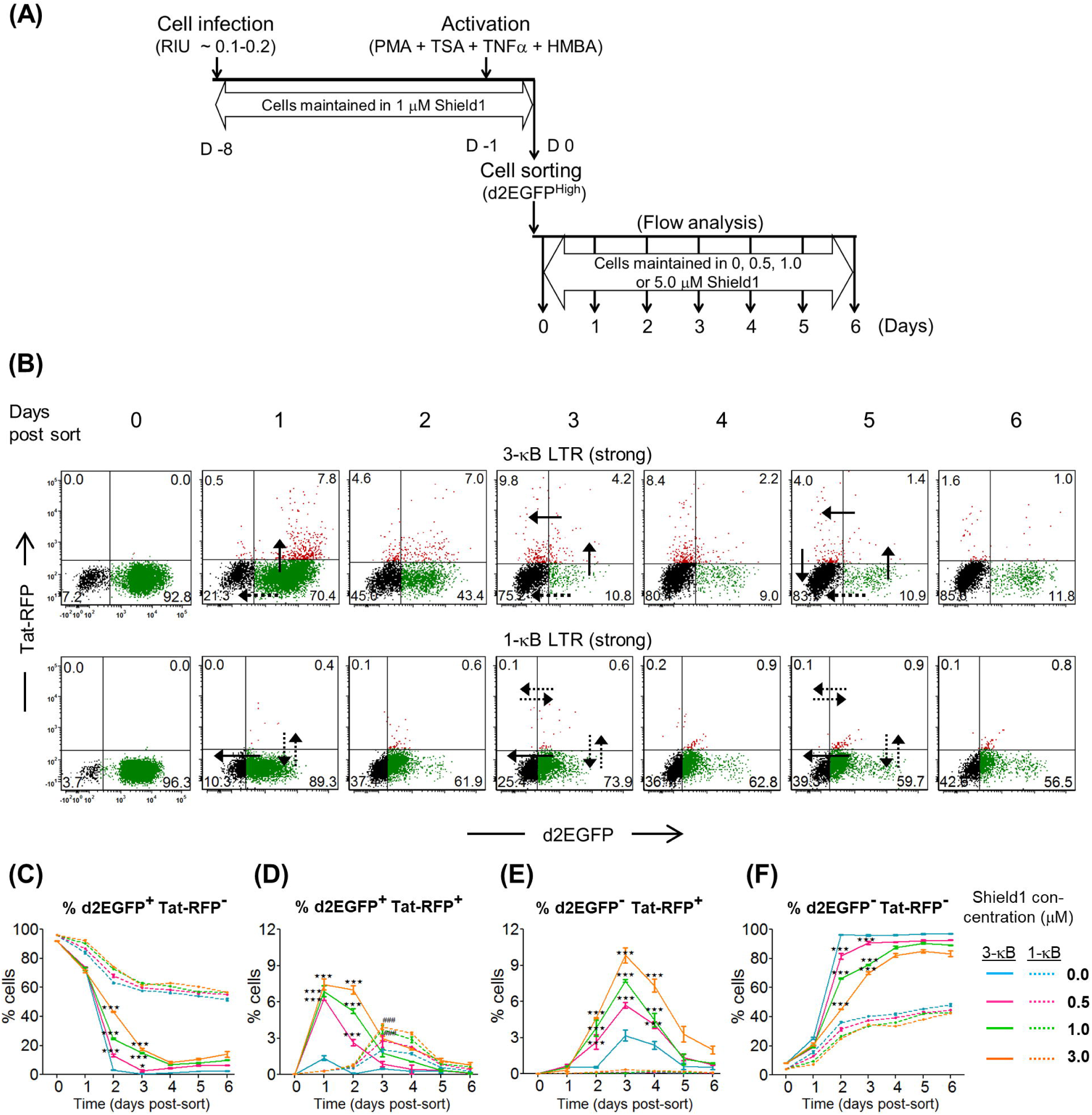
The TTF model identifies two distinct modes of latency establishment in the strong and the weak LTRs. **(A)** The experimental scheme to study latency establishment as described in Figure 4A with slight modifications. The sorted d2EGFP^High^ cells were divided into four separate fractions and maintained at different concentrations of Shield1 as depicted. The cells were analyzed every 24 h for d2EGFP and Tat-RFP expression using flow cytometry. **(B)** The temporal d2EGFP and Tat-RFP trajectories for the strong (3-κB; upper panel) and the weak (1-κB; lower panel) LTRs at the 3 μM Shield1 concentration are presented. The black-solid and the black-dotted arrows denote the dominant and the less dominant routes of transit for each LTR-variant, respectively. Individual kinetic curves of the four distinct populations- **(C)** % d2EGFP^+^ Tat-RFP^-^, **(D)** % d2EGFP^+^ Tat-RFP^+^, **(E)** % d2EGFP^-^ Tat-RFP^+^, and **(F)** % d2EGFP^-^ Tat-RFP^-^. Mean values from experimental triplicates ± SD are plotted. Data represent three independent experiments. Two-way ANOVA with Bonferroni post-test was used for the statistical evaluation (*p<0.05, ***p<0.001 and ns - non-significant). The solid- and dotted-coloured curves represent various concentrations of Shield1 for the 3-κB and the 1-κB LTRs, respectively.

The TTF model of latency offered several essential insights. Importantly, the ability to visualize two different fluorescent proteins (d2EGFP and Tat:RFP:DD) co-expressed under the LTR permitted to identify the different stages of the viral gene expression and latency, which we collectively refer to as the viral ‘latency cycle’. Although both the fluorescent proteins were expressed under the control of the same viral promoter, the expression of d2EGFP was perceptible earlier and at a higher intensity than that of the Tat:RFP:DD fusion protein. The increased molecular size of the Tat:RFP:DD fusion protein, the slow maturation of DsRed2, and the compromised translation efficiency due to the IRES element, all may have contributed to the observed difference between the d2EGFP and Tat-RFP expression profile (Figure 7B, Day 0). The profile of gene expression through the different phases of the latency cycle is remarkably different between the two viral promoters. The transiting of the cells through the successive phases of the latency cycle is illustrated explicitly when the 3-κB LTR profile is examined (Figure 7B, top panel). At Day 0 following the d2EGFP^High^ sort, the vast majority of cells (92.8%) were d2EGFP^+^ Tat-RFP^-^ representing a transcriptionally active viral promoter (Figures 7B, top panel; Day 0 and 7C). During the following 24 hours, the d2EGFP^+^ Tat-RFP^-^ cells exited this compartment via two distinct and diagonally opposite routes. While a significant proportion of these cells (approximately 15%) switched off d2EGFP expression to directly return to the d2EGFP^-^ Tat-RFP^-^ compartment, approximately 6.6% of cells up-regulated Tat-RFP expression from the 3-κB LTR to transit to the d2EGFP^+^ Tat-RFP^+^ compartment alluding to a strong Tat-dependent transcriptional activity (Figures 7B, top panel; Day 1 and 7D). At the subsequent time points, d2EGFP^+^ Tat-RFP^-^ cells continued to vacate this compartment using both the exit routes to reach the d2EGFP^-^ Tat-RFP^-^ compartment such that on Day 6, 84.3% of the viral strains re-established latency under the strong viral promoter. Importantly, the cells in the d2EGFP^+^ RFP^+^ compartment, unlike those of the d2GFP^+^ Tat-RFP^-^ compartment, appeared to move to latency only in one direction to the d2EGFP^-^ Tat-RFP^+^ compartment (Figures 7B, top panel; Day 3 and 7E). The relative proportion of the cells present in the d2EGFP^-^ Tat-RFP^+^ compartment was significantly higher than that of the d2EGFP^+^ Tat-RFP^+^ compartment at time points after Day 1 alluding to the unidirectional movement of these cells to latency. Importantly, the d2EGFP^-^ Tat-RFP^+^ compartment is unique since this quadrant represents the proviruses that have ‘recently’ switched off transcription, with significant levels of physiological Tat still persistent in the system as indicated by the RFP^+^ phenotype. The proviruses of the d2EGFP^-^ Tat-RFP^+^ compartment also transited to latency only in one direction and entered d2EGFP^-^ Tat-RFP^-^ compartment (Figure 7B, top panel; Day 5 and 7F).

In contrast, the 1-κB LTR predominantly displayed the Tat-independent transactivation (Figure 7B, bottom panel). Although approximately 4% of these cells expressed Tat-RFP at a Shield1 concentration of 3 μM, the Tat-RFP expression was delayed by 24 h, as compared to that of the 3-κB LTR, with the Tat-RFP expression reaching a peak only on D3. Importantly, despite the presence of Tat, these dual-positive cells of 1-κB LTR (d2EGFP^+^ Tat-RFP^+^) did not move forward to the d2EGFP^-^ Tat-RFP^+^ compartment, unlike those of 3-κB LTR, but returned to the d2EGFP^+^ Tat-RFP^-^ quadrant (Figure 7B, bottom panel; Day 3). The proviruses activated by Tat-independent transactivation primarily manifested the d2EGFP^+^ Tat-RFP^-^ phenotype, and these viruses returned to latency by switching off the d2EGFP expression and typically not inducing Tat-RFP expression. While a large majority of 3-κB LTR viral strains and nearly all the viral strains of 1-κB LTR followed this route of latency, a smaller proportion of proviruses of 3-κB LTR moved forward activated by Tat-dependent transactivation that manifested the d2EGFP^+^ Tat-RFP^+^ phenotype. Approximately 14% of the 3-κB LTR viral strains were activated by the Tat-dependent transactivation that followed a unidirectional trajectory to latency via the d2EGFP^+^ Tat-RFP^+^ and d2GFP^-^ Tat-RFP^+^ compartments. In contrast, approximately, only 1% of 1-κB LTR viral strains could follow the Tat-dependent transactivation, while the reminder induced only by the Tat-independent activation and returning to latency directly from the d2EGFP^+^ Tat-RFP^-^ compartment. Thus, the transcriptional strength of the viral promoter appears to play a critical role in not only regulating the activation of viral gene expression but also the latency kinetics and whether or not Tat-dependent transactivation is recruited to the LTR. Only the strong 3-κB LTR, but not the weak 1-κB LTR, could successfully undergo Tat-dependent transactivation. The major routes of entry into latency (Tat-dependent or Tat-independent) exhibited by the two LTR variants are shown in solid-black arrows, while their less dominant trajectories are indicated in dotted-black arrows in Figure 7B. Individual trajectories of the percentages of the four distinct fluorescent populations are presented (Figures 7C, 7D, 7E, and 7F).

Interestingly, a clear demarcation in the profiles of the weak and strong LTRs is evident at the level of the Tat-independent transactivation – in the Tat-RFP negative cell populations (Figures 7C and 7F). The d2EGFP^+^ Tat-RFP^-^ cells of the 3-κB LTR down-regulated d2EGFP at all the concentrations of Shield1 by D4. In contrast, latency establishment in the same population of the 1-κB LTR was incomplete, and nearly half of these cells remained d2EGFP^+^ on D6. This was primarily because a subset of the d2EGFP^+^ Tat-RFP^-^ cells at the later time points (D1 and beyond) followed the Tat-dependent route to latency in the case of the strong 3-κB, but not the weak 1-κB promoter. Therefore, from the data of the TTF model, it appears that Tat-dependent transactivation can silence the promoter at a faster rate as compared to that of the Tat-independent pathway. Further, the kinetics of percent d2EGFP^+^ to d2EGFP^-^ transition, irrespective of the Tat-RFP expression, demonstrated an identical pattern of promoter silencing in the TTF model with the strong promoter (3-κB) facilitating a faster rate of silencing compared to the weak promoter (1-κB) when compared with the ATF model (3- and 4- κB vs. 2-, 1- and 0-κB LTRs). At all the concentrations of Shield1, the strong 3-κB LTR switched off faster than the weak 1-κB LTR. Thus, the data obtained from the TTF model are strongly suggestive that the transcriptional strength of the HIV-1 LTR plays a critical role in controlling viral latency as a validation of the ATF model. A strong LTR is not only faster in establishing viral latency but also is rapid in revival kinetics from latency, whereas a weak viral promoter appears to be restricted in both the functions.

### A sustained presence of Tat in the nucleus following the LTR switch-off

The latency kinetics of two different cellular models (ATF and TTF) alluded to the direct involvement of Tat in the transcriptional suppression of the viral promoter, in a concentration-dependent manner. Furthermore, we could detect the presence of Tat-RFP fusion protein in cells harboring a transcriptionally silent provirus (d2EGFP^-^ Tat-RFP^+^) containing a strong viral promoter (3-κB-LTR) (Figure 7). Therefore, it was necessary to evaluate the physiological levels and the relative distribution of Tat in cells concomitant with LTR-silencing. To this end, we tracked the expression pattern of the Tat protein in Jurkat cells using indirect immunofluorescence while the cells transited from the ‘ON’ to the ‘OFF’ state. Jurkat cells infected with the J-cLdGIT-3-κB viral strain encoding d2EGFP (ATF model) were monitored at 4 - day intervals up to day 16 for d2EGFP expression using flow cytometry (Figure 8A, left panel). At Day 0, the d2EGFP^High^ cells were sorted and subjected to indirect immunofluorescence for Tat at different points (D0, D4, D8, D12, D14, and D16) (Figure 8A, right panel). A combination of a high-titer, polyclonal, rabbit anti-Tat primary antibody, and an anti-rabbit, Alexa-568 conjugated secondary antibody was used in the assay.

**Figure 8:**
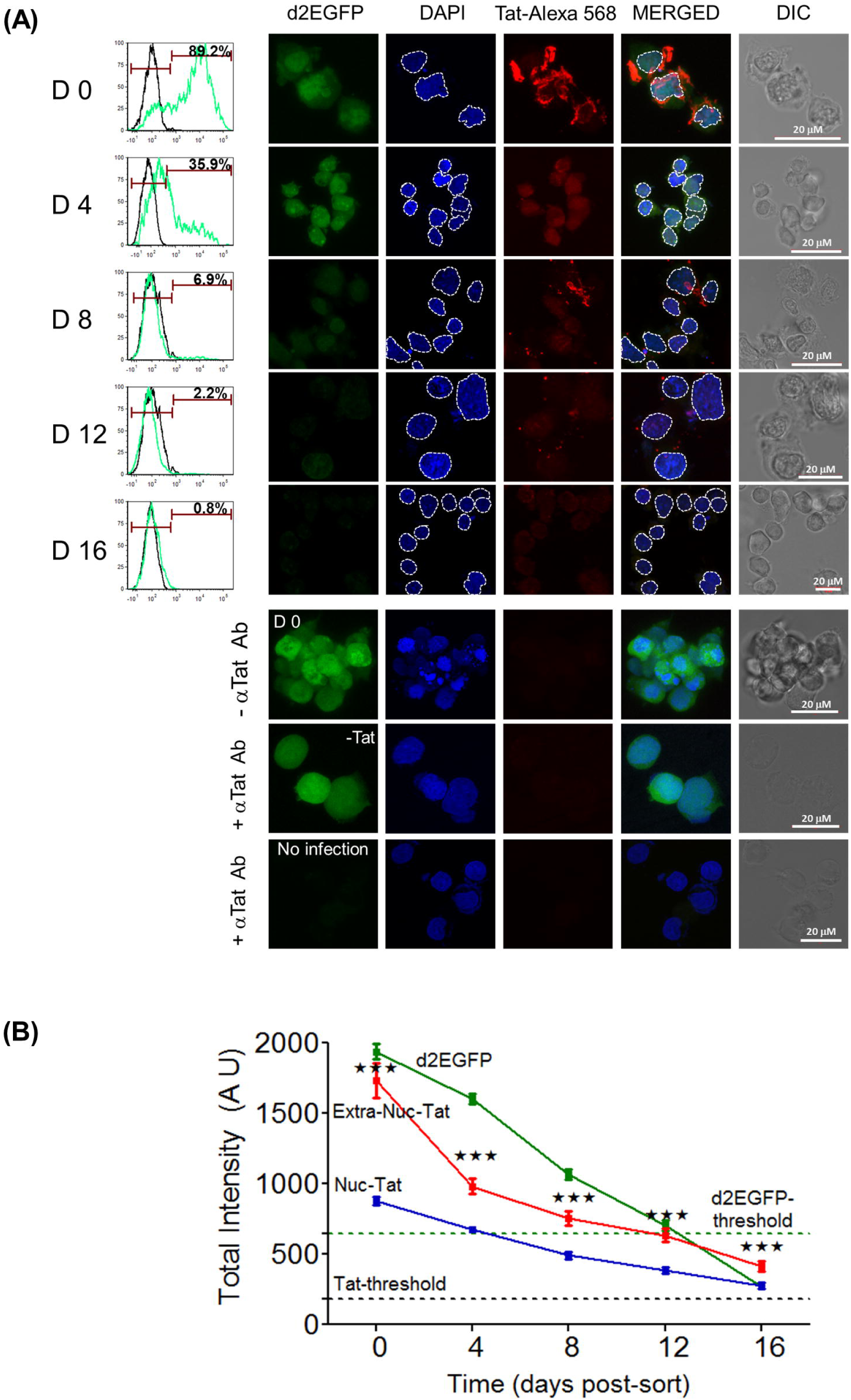
Persistent presence of Tat in latent cells. **(A)** The temporal profile of Tat expression during LTR-silencing in a stable J-LdGIT-3-κB cell pool of the ATF model. The experimental strategy of latency establishment was similar as in Figure 4A. At defined time points, a small fraction of the sorted d2EGFP^High^ cells was flow-analyzed for d2EGFP expression (left panel), while the remaining cells were subjected to indirect immunofluorescence of Tat using a rabbit anti-Tat primary antibody and an anti-rabbit Alexa- 568 conjugated secondary antibody. DAPI was used to stain the nucleus. Representative confocal images of Tat and d2EGFP-expression at indicated time points (right panel) are presented. Appropriate negative controls for Tat IF are presented in the bottom panels. The white dotted line demarcates the nucleus from the extra-nuclear compartment in each cell. Scale bar = 20 μM. **(B)** The quantitative analysis of d2EGFP and Tat-Alexa 568 intensity levels in the nuclear and extra-nuclear compartments (arbitrary units) at multiple time points. Data from 150 individual cells at each time point and three independent experiments are presented. The threshold values for total cellular d2EGFP and Tat-Alexa 568 intensities were obtained from uninfected Jurkat cells (Figure 8A; Lane-8) and infected, unstained cells (Fig 8A; Lane-6), respectively. Mean values ± SEM are plotted. One-way ANOVA was used for statistical evaluation (***p<0.001).

d2EGFP expression analysis by flow cytometry found a progressive downregulation of the fluorescence, and by D8 and D16, only 6.9% and 1% of the cells, respectively, remained positive. The profile of d2EGFP expression of individual cells captured by confocal microscopy was perfectly consistent with that of the flow analysis; and, visible fluorescence could not be detected at D8 and beyond. However, trace levels of Tat expression as determined by the indirect immunofluorescence of Tat (Tat-Alexa 568 signal) could be visually noted above the background on D12 and D16 despite the complete downregulation of d2EGFP, indicating the sustained presence of Tat in an LTR-OFF context. Of note, Tat expression was found in two different compartments of the cells, nuclear and extra-nuclear, the latter mostly localized to the cell membrane. Fluorescent intensities of d2EGFP expression, as well as that of Tat-Alexa 568, were determined independently in the nuclear and the extra-nuclear compartments of 150 individual cells, at all the time points (Figure 8B). The threshold levels of fluorescent protein expression were determined by using uninfected Jurkat cell control for d2EGFP (Lane-8; n = 10) and no-primary antibody control for Tat (Lane-6; n = 10). Importantly, while the fluorescence of d2EGFP reduced progressively with time and fell below the threshold by D12 representing the establishment of latency, the fluorescence of Tat, in either compartment, did not drop below the Tat-Alexa 568 threshold even at D16. The slopes of reduction of the Tat intensities during the initial phases (D0 to D4) of latency-establishment were estimated to be −74.54 ± 16.8 and −37.28 ± 3.2 in the extra-nuclear and nuclear compartments, respectively. At the later time points (D8, D12, and D16), there was only a moderate reduction in the Tat levels in either of the compartments. The data are thus suggestive of a higher level of stability of Tat in the nucleus with possible implications for HIV latency. Importantly, the data of Tat-immunofluorescence are in perfect agreement with the results of the TTF model, where the few d2EGFP^-^ Tat-RFP^+^ cells at the later stages of promoter-silencing indicated sustained presence of low-levels of Tat molecules in the LTR-switched OFF cells (Figure 7B; top panel). In summary, immunofluorescence not only detected the presence of Tat in the latently infected cells as late as D16 post-sorting but also demonstrated a rapid loss of Tat from the extra-nuclear compartment while its relative stability in the nucleus.

### The in situ proximity ligation assay (PLA) detects the presence of Tat in the latently infected cells

In indirect immunofluorescence, the over-all intensity of the Tat at D12, D14, and D16 in both the cellular compartments was only marginally above the background level. To increase the sensitivity and detect limited quantities of Tat in the ‘LTR OFF’ cells, we used the highly sensitive proximity ligation assay (PLA), which conjugates immunostaining with the rolling-circle replication and outperforms the traditional immune assays in sensitivity to detect trace amounts of endogenous proteins (30, 31). We optimized Tat-PLA in HEK293T cells using a pair of anti-Tat primary antibodies raised in different hosts (rabbit and mouse). Since PLA does not work well in non-adherent cells, and our attempts to adapt the protocol to the Jurkat cells were not successful, we used HEK293/HEK293T cells in this assay. Using sub-genomic viral vectors encoding Tat representing HIV-1B (pLGIT) or HIV-1C (pcLGIT), we optimized PLA (Figures 9A and 9B). The B-Tat protein could be detected as distinct white dots as opposed to sparse dots in no-antibody and single antibody controls (Figure 9A). Moreover, a dose-response in the intensity of PLA dots and plasmid concentration, as well as a good correlation between the PLA dot number and GFP MFI, are evident in the case C-Tat (Figure 9B).

**Figure 9:**
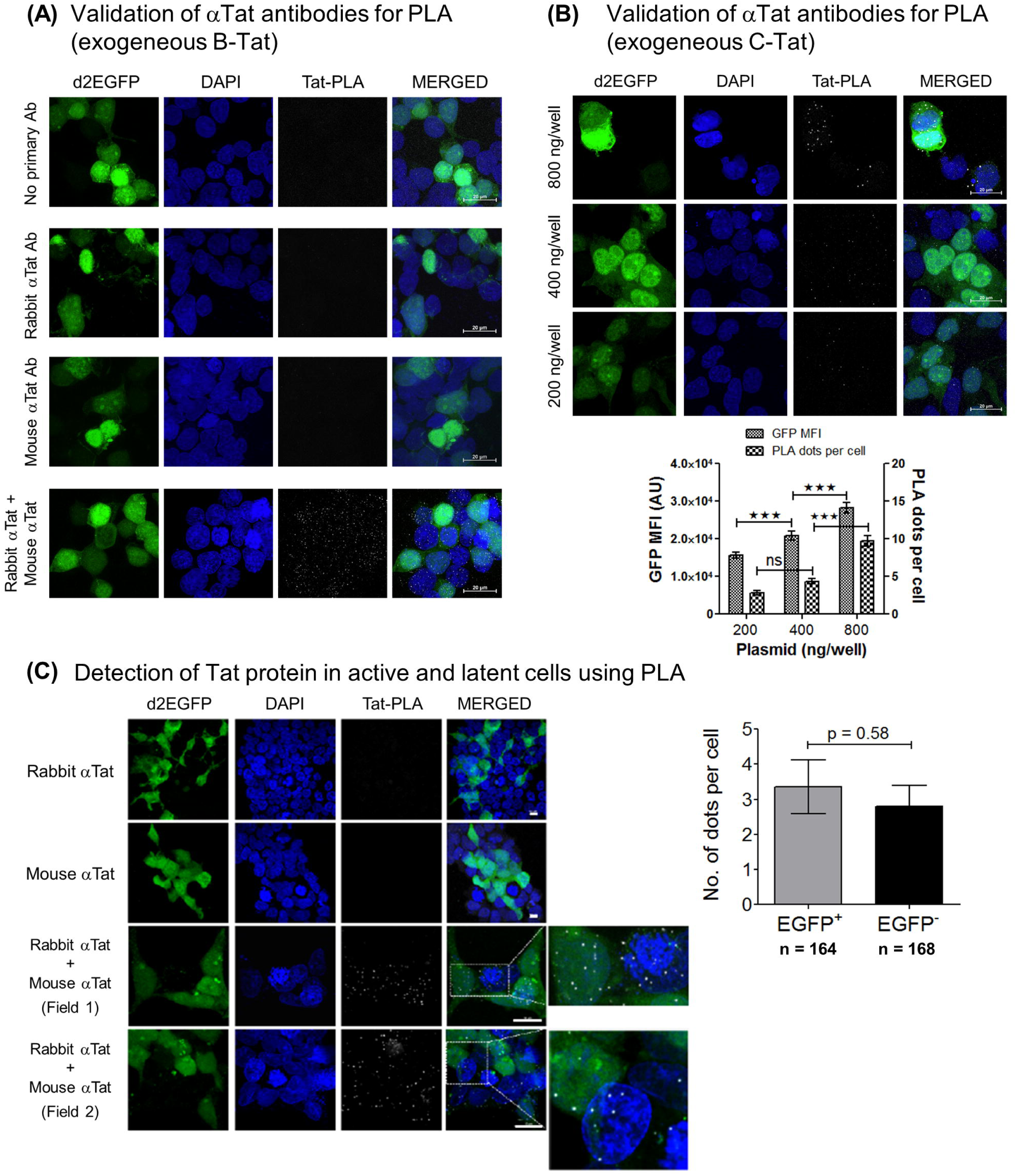
The presence of Tat in latent cells as confirmed by the highly sensitive proximity ligation assay (PLA). **(A)** The detection of exogeneous B-Tat in PLA. Approximately 0.5 million HEK293T cells seeded per well in a 12-well culture dish were transfected with 800 ng of pLGIT, an expression vector encoding Tat of HIV-1B, on poly-L-lysine coated coverslips. Tat-PLA was performed according to the manufacturer’s protocol using a pair of anti-Tat primary antibodies (rabbit-polyclonal; # ab43014, Abcam and mouse-monoclonal; # 7377, NIH-AIDS reagents program). Representative confocal images of a ‘no antibody’ control (Lane 1), single anti-Tat antibody controls (Lanes 2 and 3) and both the antibodies (Lane 4) are presented. The mean values from three independent experiments ± SEM are plotted. A one-way ANOVA with Bonferroni’s multiple comparison post-test was used for statistical analyses (***p<0.001). **(B)** A DNA dose-response of PLA using pcLGIT, an expression vector encoding Tat of HIV-1C (200, 400, 800 ng). The mean GFP intensities and the average number of PLA dots per cell for the amount of pcLGIT vector transfected are presented. **(C)** The Tat PLA dot quantitation in the active vs. latent cells was performed in HEK293 cells independently and stably infected with the cLdGIT-4-κB and 3-κB strains of the ATF panel (Figure 2A). Cells were infected with one of the viral strains (∼ 0.5 RIU), d2EGFP^High^ cells were sorted, the cells were incubated for proviral-LTR relaxation to arrive at a mixed population of d2EGFP^+^ (active) and d2EGFP^-^ (latent) cells, and both the cell populations were subjected to Tat-PLA. Approximately 50,000 mixed d2EGFP cells seeded in a well of an 8-well slide chamber were subjected to PLA. Representative confocal images depicting single antibody controls (Lanes-1 and 2) and Tat-PLA with both the antibodies (Lanes-3 and 4) are shown (left panel). Two sub-fields with distinct Tat-PLA dots (white) in both d2EGFP^+^ and d2GFP^-^ cells have been enlarged for clarity. The number of Tat-PLA dots per cell was determined independently for d2EGFP^+^ as well as d2EGFP^-^ cells, and the mean values from three independent experiments ± SEM are plotted. The total number of cells counted for d2EGFP^+^ phenotype was 164 (128 for 4-κB and 36 for 3-κB) and for d2EGFP^-^ phenotype was 168 (123 for 4-κB and 45 for 3-κB). A two-tailed, unpaired *t*-test was used for statistical analyses.

Using the optimized PLA protocol for Tat, we asked if the Tat protein could be detected in d2EGFP OFF cells. To this end, HEK293 cells stably and independently infected with the 4- and the 3-κB variants of the ATF-cLdGIT panel were sorted for the d2EGFP^High^ cells. After a week of incubation following the enrichment, approximately 50% of the cells expressed d2EGFP, and the cell pool contained both active (d2EGFP^+^) and latent (d2EGFP^-^) cell clusters. Tat-PLA was then performed using the mixed d2EGFP pool corresponding to both the strong LTR (3- and 4-κB) variants to quantitate the Tat-PLA signals in the alternate phenotypes (active and latent). The cells stained with either of the antibodies alone did not show any Tat-specific signals confirming the specificity of the assay (Figure 9C left panel, top two lanes). Tat-specific staining was evident only in the presence of both the antibodies not only in the d2EGFP^+^ cells but also in the d2EGFP^-^ cells (Figure 9C left panel; bottom two lanes). The average number of Tat-PLA dots per cell was determined in a total of 164 d2GFP^+^ cells (128 and 36 cells for 4-κB and 3-κB variants, respectively) and 168 d2EGFP^-^ cells (123 and 45 cells for 4-κB and 3-κB variants, respectively) comprising of three independent experiments (Figure 9C; right panel). These values were found to be 3.35 ± 0.77 and 2.8 ± 0.59 for d2EGFP^+^ and d2EGFP^-^ cells, respectively, although the difference was not significant statistically. The Tat-PLA data in HEK293 cells confirmed the presence of Tat in the latent cells at a concentration comparable to that of active viral transcription.

### Differential occupancy of cellular complexes on active and silent LTRs of bimodal clones

Burnett JC et al. examined the differential occupancy of NF-κB factors (p50 and p65) at each of the two identical NF-κB motifs (I and II) in HIV-1B LTR by introducing inactivation mutations into each of these sites individually and the corresponding impact on the transcriptional activity (26). A similar examination at the C-LTR has not been performed. A previous report from our laboratory demonstrated that NFAT1 and 2 proteins could be recruited to the C-κB motif, the variant NF-κB motif unique for HIV-1C, with an affinity superior to that of the canonical H-κB site (32). We attempted to compare the identity of the transcription factors and other host factors binding to the viral promoter between the active and suppressed states under identical experimental conditions.

Having demonstrated the presence of Tat in the cells containing the latent provirus in both the TTF (Figure 7), and ATF (Figures 8 and 9) models by flow cytometry and confocal imaging, respectively, we, next asked if Tat in these cells is recruited to the latent viral promoter. We sorted the d2EGFP^High^ and d2EGFP^-^ cell populations from the two clonal cell populations and using the chromatin immunoprecipitation assay, we examined for the presence of several essential host factors or epigenetic marks (Rel family members- p50, p65; NFAT1 and NFAT2; RNA polymerase Ser2 phosphorylation, and Histone 3 lysine 9 trimethylation), as well as Tat in the chromatin preparations of the active and latent cells. The ChIP assays were performed by amplifying a 240 bp fragment spanning the enhancer-core promoter region in the LTR using a semi-quantitative PCR and also using an independent Taqman probe-based real-time PCR amplifying a 127 bp region spanning the NF-κB and Sp1 sites in the LTR (Figure 10A).

**Figure 10:**
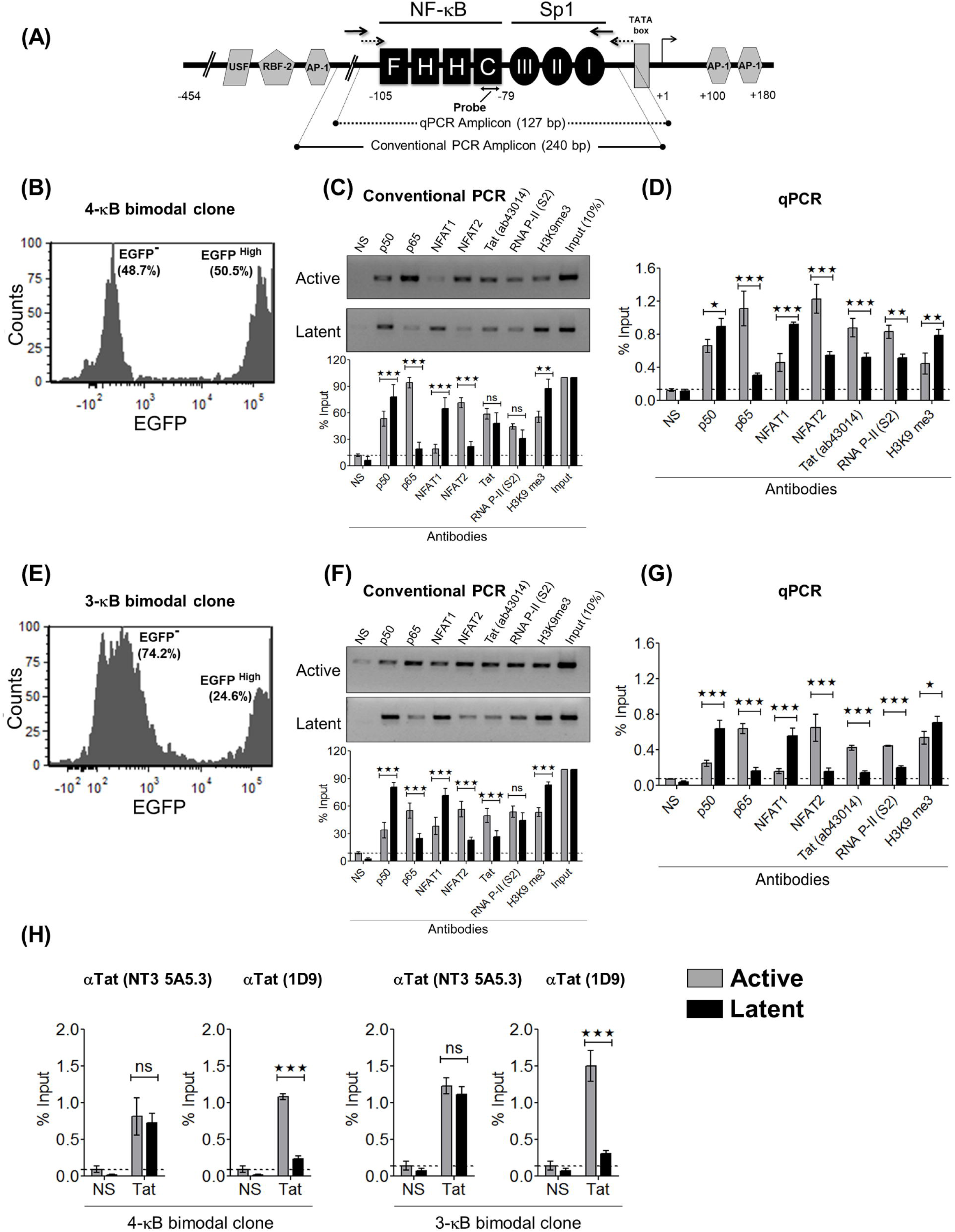
The active and latent LTRs recruit host-factors differentially. **(A)** Schematic representation of the LTR sequence and the positions of the primer pairs used in the conventional PCR (240 bp, solid-black) and qPCR (127 bp, dotted black) for the ChIP analysis is depicted. **(B)** and **(E)** The pre-sort percentages of the EGFP^-^ and EGFP^High^ cells corresponding to the 4-κB (Panel-B) and the 3-κB (Panel-E) bimodal clones are presented. **(C)** and **(F)** Data from a conventional PCR-based ChIP assay of the active (EGFP^High^) and latent (EGFP^-^) promoters for several host factors and Tat protein are presented for the 4-κB (Panel-C) and 3-κB (Panel-F) clones, respectively. Cell-lysate from two million cells (active or latent) and 2 μg of the respective antibody were used for the individual IP reactions. A rabbit polyclonal anti-Tat antibody (# ab43014, Abcam) was used for the Tat-IP. One-tenth of the IP chromatin was used as the input control. Conventional PCR was repeated thrice for each IP reaction. Representative gel image and the corresponding densitometry analyses are presented. Data for each band are normalized to the input. The mean values from quadruplicates ± SD are plotted. A two-tailed, unpaired *t*-test was used for statistical analyses (*p<0.05, **p<0.01, ***p<0.001, and ns- non-significant). A qPCR-based ChIP assay was performed independently using identical experimental conditions and **(D)** the 4-κB or **(G)** the 3-κB clones. The data for each IP was calculated using the percent-input method. The mean values from qPCR triplicates ± SD are plotted. A two-tailed, unpaired *t*-test was used for statistical analyses (*p<0.05, **p<0.01, ***p<0.001, and ns- non-significant). **(H)** Two additional qPCR-based ChIP assays using two different mouse monoclonal anti-Tat antibodies (1D9 and NT3 5A5.3) were performed using the active and latent fractions of the two bimodal clones. The mean values from qPCR triplicates ± SD are plotted. A two-tailed, unpaired *t*-test was used for statistical analyses (*p<0.05, **p<0.01, ***p<0.001, and ns- non-significant).

The comparative analysis of the nature of the host factors recruited between the active and latent promoters was highly reproducible and consistent between the 4- and 3-κB LTRs (Figures 10C, 10D, 10F, and 10G). While the transcription promoting host factors, p65 and NFAT2, and epigenetic marks RNA Pol II S2, were found associated with the active viral promoters at significantly higher levels, the transcription repressive factors, p50, NFAT1, and epigenetic marks H3K9Me3 were preferentially associated with the latent viral promoters. That the p50-p65 heterodimer is transcription-promoting, the presence of a significantly higher concentration of p65 at the active promoter is expected (33–35). On the other hand, the preferential association of p50 with the latent promoter is suggestive of the formation of the p50 homodimer, a known transcription suppressor (36). Similarly, our data are also in agreement with the previous reports regarding the transcription suppressive and supportive functions of NFAT1 and NFAT2, respectively (37–39).

The most crucial finding of the present study is the detection of the association of the Tat protein with the latent LTR. The Tat protein was found associated with the latent 4-κB and 3-κB promoters at levels 1.7- and 3-folds lower, respectively, as compared to their active counterparts. The results were reproducible and consistent in the semi-quantitative PCR-based ChIP analyses between the two strong viral promoters (Figures 10C and 10F). The data were also consistent between the conventional and the quantitative real-time PCRs performed following immunoprecipitation (Figures 10D and 10G). To the best of our knowledge, the present study is the first one to demonstrate the association of Tat with the latent LTR. The above ChIP data were generated using a commercial rabbit polyclonal anti-Tat antibody (Cat # ab43014, Abcam). The data were also reproducible when two additional mouse monoclonal anti-Tat antibodies targeting different epitopes in Tat (Cat # 7377 and # 4374, NIH AIDS reagent program, Maryland, USA) were used in the assay (Figure 10H). All the three different anti-Tat antibodies furnished positive ChIP signals for Tat at both the latent viral promoters (3- and 4-κB), over and above the respective IgG-isotype controls.

## Discussion

### The significance of Tat recruitment to the latent LTR

The primary finding of the present work is the identification of a positive correlation between the transcriptional strength of the LTR and faster latency kinetics via the mediation of proportionately enhanced Tat concentration. We found that at the time of commitment towards latency and at subsequent time points, the intracellular concentration of Tat is not a limiting factor, thus, ruling out the possibility that the limiting levels of Tat underlie latency establishment. To exert a positive or negative influence on the LTR, Tat must be present in the nucleus and recruited to the viral promoter. Using three different experimental strategies, the flow analysis of the Tat-RFP fusion protein (Figure 7), indirect immunofluorescence (Figure 8, and a proximity ligation assay (Figure 9C), we successfully demonstrated the presence of Tat in the nucleus of the latent cell, through the successive stages of latency establishment. The presence of Tat could be detected in both the nuclear and cytoplasmic compartments by confocal microscopy. Additionally, using ChIP, we could ascertain the recruitment of Tat to the latent LTR using three different anti-Tat antibodies, although Tat levels in the latent nuclei were typically inferior to those of the active nuclei (Figure 10). Furthermore, a weak but discernible signal of the Tat-transcripts was evident in the latent fractions of the bimodal clones of both the strong LTRs used in the assay (Figures 5F and 5J). All these data are strongly suggestive that Tat plays a direct role in promoting latency. Unfortunately, our attempts at adopting PLA to suspension cells were not successful. How Tat is recruited to the chromatin complex needs to be determined. A few studies previously showed the direct binding of Tat to extrachromosomal HIV-1 promoter and proviral DNA (40, 41); however, the tethering of Tat to the nascent TAR element, as a part of the paused RNA PolII complex proximal to the latent promoter (42), is a more likely possibility.

### The underlying mechanisms regulating HIV-1 latency remain enigmatic

There have been several attempts to understand HIV-1 latency as this question contains direct relevance for clinical management and viral purging (43). The complexity of HIV-1 latency has led to two distinct schools of thought to explain the phenomenon - the hypothesis of ‘epiphenomenon’ where the host environmental factors including the epigenetic modifications play the deterministic role (1, 44), and that of ‘viral circuitry’ where decision making is hardwired in the intrinsic Tat-LTR regulatory circuit (7, 9). The two models, which need not necessarily be mutually exclusive, have been supported by considerable experimental evidence but also have specific limitations.

Experimental evidence for epigenetic modifications controlling HIV-1 latency is available from studies using clonal cell populations typically harboring sub-genomic viral reporter vectors (45). The major limitation of this experimental model is the prolonged periods required for the cells to establish latency. The majority of individual clonal populations reach 50% of latency on an average in 30 to 80 days, which probably is not representative of the kinetics of natural latency. Given the cytotoxic properties of the viral products and the immune response, and a relatively short life-span of infected T cells (∼ 2.2 days) (46), viral gene expression is expected to drive viral evolution towards rapid, not prolonged, latency establishment, in natural infection. Additionally, it is also not understood how epigenetic silencing of an active viral promoter is ever achieved, especially in the presence of abundant quantities of Tat.

The contrasting model explaining HIV-1 latency based on the intrinsic and virus-driven stochastic phenomenon is also supported by compelling experimental evidence (7, 9). The ‘feedback-resistor’ module (12), considers a single type of chemical modification, acetylation, and deacetylation, of Tat, serving as the ‘resistor’ or dissipater of the positive transcription loop to ensure a stable latent state. The model doesn’t take into account several other PTMs of Tat. Whereas di-methylation of the lysine residues at positions 50 and 51 (47) and the arginine residues at positions 52 and 53 (48) can suppress Tat-transactivation, mono-methylation of the lysine residues shows the opposite effect (49). Importantly, methylated Tat is expected to have enhanced cellular stability with implications for latency (50). In addition to acetylation, the phosphorylation of multiple serine and threonine residues can cooperatively enhance Tat transactivation (51, 52). Polyubiquitination of Tat can also enhance the stability of Tat, thereby augmenting its transactivation function (53). Apart from the chemical modifications, Tat is also known to be inactivated by the propensity of the protein to make dimer and multimer forms, although experimental evidence is scanty in this regard (54). Furthermore, the differential forward and reverse reaction kinetics of Tat acetylation have been evaluated only in HeLa cells, but not in cells of physiological relevance to HIV-1 infection (55). Weinberger et al., also analyzed the feedback strength in terms of the noise autocorrelation function and demonstrated that a stronger Tat feedback would yield transcriptional pulses of longer durations leading to cell lysis while the weaker Tat feedback and the Tat-independent transcription would generate shorter transcriptional pulses leading to latency (8). The model, however, doesn’t reconcile to the fact that a small proportion of T cells can still escape cell death following Tat-mediated transcription and establish a viral reservoir (56). In summary, despite significant experimental evidence, the question regarding the critical deterministic factor(s) regulating HIV-1 latency remains unresolved.

### Subtype-associated molecular features may offer vital clues to HIV-1 latency

Although the fundamental constitution of the HIV-1 promoter is highly conserved among the various genetic subtypes of HIV-1, there exist many subtype-specific molecular features that may modulate gene expression considerably. Such differences are evident in the copy-number and nucleotide sequences of different TFBS especially those of USF, c-Myb, LEF-1, Ets1, NF-AT, Ap-1, NF-κB, and Sp1 binding sites, and regulatory elements such as the TATA box and the TAR element (57–59). These regulatory elements play critical roles in positively regulating the basal and inducible levels of viral transcription (60, 61). Most of the TFBS, especially the AP-1, USF, NFAT, NF-κB, and Sp-1 motifs also play a critical role in regulating viral latency by recruiting chromatin-modifying complexes and transcription suppressing factors such as the histone deacetylases (HDACs) to the viral promoter (62–66).

Of the various TFBS, both NF-κB and Sp-1 motifs are represented by multiple and tandem binding sites in the LTR, and, play crucial roles in regulating gene expression and latency (36, 67–69). The most striking feature in the HIV-1C LTR is the copy-number difference of NF-κB motifs, the sequence variation of the additional κB motifs (25), and the associated sequence variation of the Sp1III site (32). We demonstrated previously that NF-κB site duplication is unique for HIV-1C not recapitulated by any other HIV-1 genetic family. Importantly, in HIV-1C, a unique NF-κB motif (the C-κB element, GGG<u>G</u>C<u>G</u>TTCC) and a genetically distinct and subtype-specific Sp1III site are located at the core of the promoter, and the two elements establish a functional association in enhancing HIV-1C transcription (32).

HIV-1C could serve as an ideal model to ask whether transcriptional strength can affect viral latency, a property not explored previously probably due to the absence of NF-κB copy-number variation in non-HIV-1C subtypes. The rapid expansion of the 4-κB viral strains in India in a short period of ten years, from 2% to 25 – 35%, is quite surprising (25). The ATF model we used here also demonstrated a perfect positive correlation between the number of NF-κB binding sites in the LTR and the viral transcriptional output in the form of EGFP and Tat transcripts (Figure 2) suggesting that all the four NF-κB binding sites in the LTR are functional. In this backdrop, it remains intriguing why HIV-1C strains require enhanced transcriptional strength, and despite having a strong promoter, how do these viral strains establish and maintain latency while the other HIV-1 genetic subtypes do not adopt such an evolutionary strategy.

### Reciprocal binding of host-factors at the active and latent promoters

Gene expression is the outcome of multiple layers of regulatory events consisting of the *cis*-acting TFBS and the *trans*-acting chromatin remodelers, viral factors, especially Tat, epigenetic marks, and a cross-talk between a wide array of proteins, ultimately leading to diverse phenotypic outcomes. Numerous studies attempted to examine how the nature of the host factor complexes recruited at the LTR regulates the dynamic switching between the active and latent states (45, 70). In an elegant analysis, Burnett et al., used PheB cells derived from the GFP^Mid^ parental Jurkat cells (analogous to GFP^Dim^ in (7) and GFP^Low^ in the present study) and compared the nature of cellular complexes recruited between transcriptionally active and latent cells (26). This study demonstrated a non-overlapping function of the two genetically identical NF-κB sites in regulating transcriptional activation versus suppression. We employed a similar experimental strategy with the exception that we used the EGFP^High^ clonal cell populations (the bimodallers, Figure 5B) that manifested a bimodal phenotype of EGFP expression. The EGFP^High^ and the EGFP^-^subsets of bimodal cell clones offered a significant technical advantage of normalizing the inherent differences in cellular parameters.

The preferential binding of p50 and p65 (RelA) at the latent and active promoters ascertained the repressive and inducing functions of p50-p50 homodimer and p50-RelA heterodimer respectively, of HIV-1 transcription (Figure 10) (26, 35, 36). Unlike the NF-κB proteins, the impact of individual NFAT members on HIV-1 latency has not been examined in great detail. To the best of the knowledge, the present study is the first to demonstrate a reciprocal binding pattern of NFAT1 and NFAT2 at the active and latent promoters, respectively, in the context of clonal cells. Since NF-κB and NFAT factors share overlapping sites (71–73), NFAT may have a significant influence on latency in HIV-1C. Furthermore, the NF-κB sites in the C-LTR (F, H, and C-κB sites) are genetically different, adding to the multitude of possible combinations. Targeted inactivation of each κB site, one at a time, followed by ChIP, may provide meaningful insights into the contribution of each κB sequence to diverse signaling pathways and HIV-1C latency.

The key finding of the present study, however, is the detection of the association of the Tat protein with the latent LTR. The results were highly reproducible and consistent between the two strong viral promoters (Figure 10). The data were also consistent between the conventional PCR and the quantitative real-time PCR performed following immunoprecipitation. The data were reproducible when three different anti-Tat antibodies targeting different epitopes in Tat were used in the assay. The Tat protein was found associated with the active 4-κB and 3-κB promoters at 1.7- and 3-folds higher, respectively, as compared to their latent counterparts. To the best of our knowledge, the present study is the first one to demonstrate the association of Tat with the latent LTR, albeit at a lower intensity as compared to the active promoter.

### The negative feedback circuits represent a powerful and common strategy biological systems exploit to regulate gene expression

Negative feedback circuits can rapidly switch off signaling cascades; therefore, this mode of gene regulation represents the most common strategy biological systems exploit to regulate gene expression (74). Molecules of biological significance controlling powerful signaling cascades such as cytokines and transcription factors, often attenuate their own production using negative feedback loops. The transcription factor NF-κB that controls the expression of numerous cellular factors that regulate a wide variety of cellular processes, down-regulates self-expression by activating the inhibitor protein IκBα (75). Likewise, interleukin-2 (IL-2), the most potent cytokine that regulates T cell viability and proliferation, limits self-production by activating the expression of a FOXP3-mediated negative feedback loop (76). Given that the latency establishment is central for HIV-1 survival towards evading immune surveillance and minimizing cytotoxicity, an active molecular mechanism would be necessary to suppress gene expression from the LTR rapidly. The decision making to achieve such a critical phase of the viral life cycle must be an intrinsic characteristic of the MTRC of the virus; it couldn’t be left to stochastic phenomena or epiphenomena regulated by cellular events. That the MTRC of HIV-1 comprises of only two elements – the LTR and Tat, and that the latter is the only factor encoded by the virus, Tat is the viral factor best positioned to regulate viral transcription and transcriptional silence both, perhaps at different phases of the viral life cycle following integration. Data presented here using two different latency cell models are not only consistent with this critical biological function ascribed to Tat but also provide additional information on latency. In the present work, we examined the latency profile only in the context of HIV-1C, and its validity must be examined in other genetic families of HIV-1.

Based on several facts, the master regulator of the virus is well-positioned to be a potential candidate to impose negative feedback on the LTR, in a temporal fashion; including the absence of a known transcription suppressor encoded by the virus, the ability of Tat to constitute the master regulatory circuit of the virus in combination with the LTR in the absence of other viral factors, the presence of Tat in the latent cell detected reproducibly and also recruited to the latent promoter, and the identification of a positive correlation between the transcriptional strength of the LTR and the rate of latency establishment.

### A Tat-dependent negative feedback mechanism to establish latency?

Based on the present study, we propose a novel model for the transcriptional repression of HIV-1 through a Tat-negative feedback mechanism. The attenuation of Tat-positive feedback signaling has been proposed to cause the LTR silencing, triggered by extracellular cues (deterministic model) or limiting Tat levels probabilistically (stochastic model) (7, 8, 12, 26). In either case, Tat concentration gradually falls below a threshold insufficient for self-renewal or successful transcriptional elongation.

Our data allude to a concentration-dependent inter-conversion of the active form of Tat to a repressive form, the latter competing with the former, strengthening a negative-feedback circuit leading to the rapid silencing of the promoter (Figure 11). We propose that the autonomous Tat-feedback loop initially favors the steady accumulation of Tat molecules to enhance transcription. Subsequently, at a point when Tat intracellular concentration surpasses a specific threshold level, Tat switches to the suppression mode down-regulating transcription, possibly depending on differential PTM modifications of Tat itself. Hence, our model proposes that the strong promoters (3- and 4-kB LTRs) characterized by a stronger Tat-feedback, can initiate a rapid transcriptional silence as compared to the weak promoters (2-, 1- and 0-κB LTRs).

**Figure 11:**
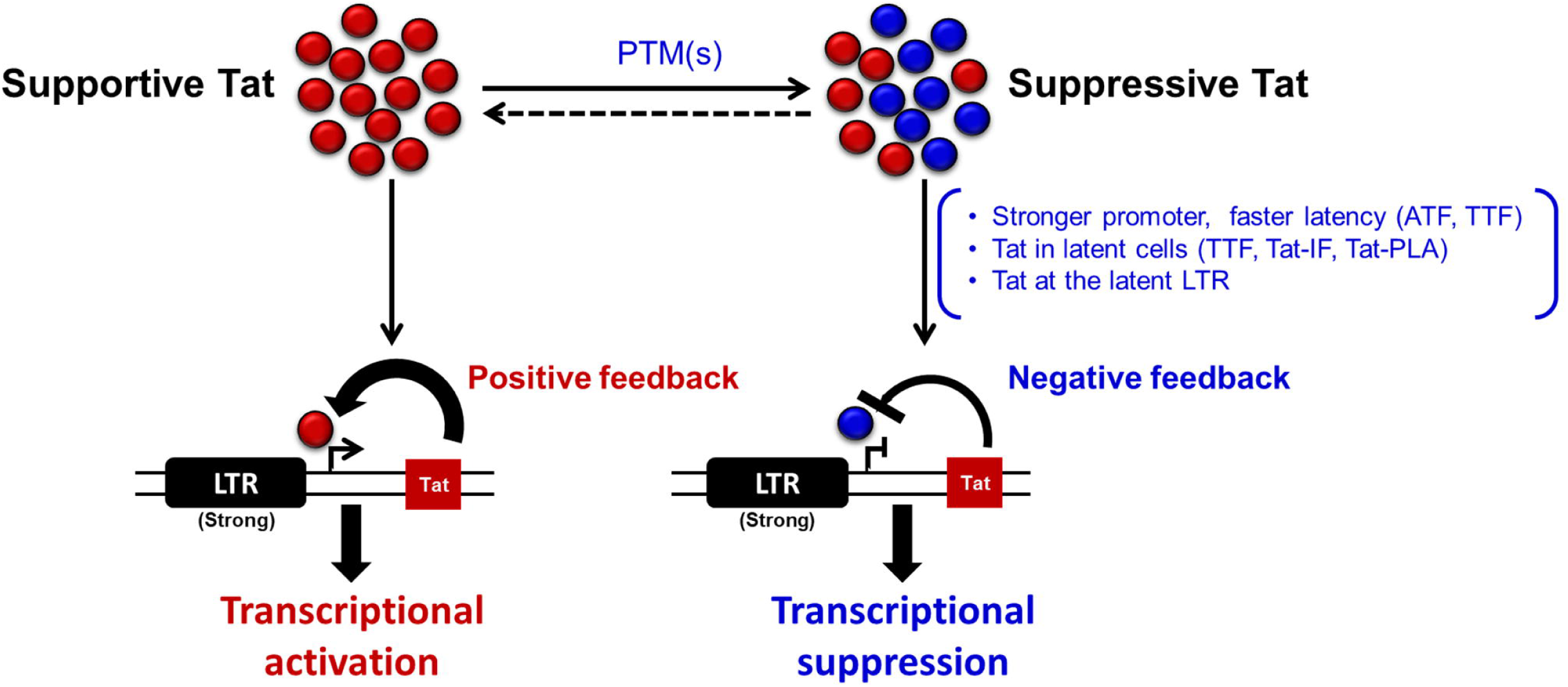
A hypothetical model depicting a concentration-dependent switch of the Tat protein through differential PTM(s) that mutually serve as an activator or suppressor at different phases of viral transcription. The model suggests that the diverse PTM(s) of Tat play a critical role in permitting the transactivator to toggle between being an activator or suppressor of transactivation of the LTR. At the time of commitment to latency, the intracellular concentrations of Tat are not limiting. Tat may initiate a negative feedback response of viral transcription in a concentration-dependent manner by regulating the expression of host factors that control the PTM(s) of Tat. The negative feedback effect of Tat, therefore, follows its initial positive feedback on the viral promoter. A viral promoter characterized by stronger transcriptional activity, thus, mediates the establishment of latency at a faster rate by producing more quantities of Tat. The data presented in this work are consistent with this model.

Our data raises several important questions related to HIV-1C latency, which were beyond the scope of the present study. Is the LTR of HIV-1C likely to continue to acquire additional copies of NF-κB and/or other transcription factor binding sites to augment transcriptional strength further? Of note, unpublished data from our laboratory (Bhange D et al, unpublished data) demonstrate a recent trend of emergence of at least 10 different types of TFBS variant HIV strains in India. Further, how the variant NF-κB motifs unique for HIV-1C modulate viral latency? Answers to these questions will shed light on the mechanism of HIV-1 latency and likely to help design novel therapeutic strategies to purge HIV infection.

We acknowledge a few technical limitations of the present work that were beyond our control. Our attempts to extend the observations to full-length HIV-1C molecular clones were not successful for two main reasons- (1) the lack of good molecular clones representing HIV- 1C (only four molecular clones of HIV-1C are available). (2) Additionally, unlike NL4-3, an HIV-1B molecular clone mostly used in the field, the few available HIV-1C clones do not lend themselves for significant molecular manipulation of any kind; the in vitro infection property of the HIV-1C viral strains is significantly low compared to other subtypes (please see a recent review (77), which highlights this problem of HIV-1C explicitly), Further, the engineering of a fluorescent protein as a reporter will make these clones nearly unviable. Our data were derived from Jurkat cell infection- the most popular cell model in the field (26, 45, 78–81). Latently infected primary cells in vivo are extremely rare in the order of 10^-6^ (1), and it would be a daunting task to isolate Tat from such a small population. Further, to examine latency establishment in primary cells, a stable cell pool must be generated, reactivated. Such manipulations in primary cells would require at least 6 to 8 weeks, a time frame not conducive to sustaining primary cells in culture. Primary cell models have traditionally been used to examine latency reversal and to evaluate latency-reversing agents but rarely to study latency establishment. A suitable, long-lasting primary cell-based model to study the mechanisms involving HIV-1 latency establishment is an absolute requirement, and we have been optimizing this model in our laboratory for future studies. Additionally, we have been raising antibodies specific to defined PTM of Tat to investigate how Tat-PTM may modulate HIV-1 latency.

## Materials and Methods

### Cell culture

Jurkat cells were maintained in RPMI 1640 medium (R4130, Sigma-Aldrich, St. Louis, USA) supplemented with 10% fetal bovine serum (RM10435, HiMedia Laboratories, Mumbai, India), 2 mM glutamine (G8540, Sigma-Aldrich), 100 units/ml penicillin G (P3032, Sigma-Aldrich) and 100 g/ml streptomycin (S9137, Sigma-Aldrich).

The human embryonic kidney cell lines HEK293 and HEK293T were cultured in Dulbecco’s modified Eagle’s medium (D1152, Sigma-Aldrich) supplemented with 10% FBS. All the cells were incubated at 37°C in the presence of 5% CO_2_.

### Design and construction of HIV-1C reporter vector panels

#### Autonomous Tat-feedback (ATF) model

The pLGIT reporter vector (HIV-1 LTR-EGFP-IRES-Tat; (7)) was a kind gift from Dr. David Schaffer (University of California, USA), in which the two different elements, the 3’LTR, and Tat, were of HIV-1B origin (NL4-3). We substituted these two elements with analogous counterparts of HIV-1C origin (Indie_C1-Genbank accession number AB023804) and referred to the vector as pcLGIT (cLTR-EGFP-IRES-cTat; (32)). Using the pcLGIT backbone, we constructed a panel of five reporter vectors containing varying copies of functional NF-κB motifs, ranging from 0 to 4 (the p911a vector series). First, an LTR containing four tandem NF-κB motifs (FHHC-LTR; H-GGGACTTTCC, C-GGG<u>G</u>C<u>G</u>TTCC, F-GGGACTTTC<u>T</u>; variations among the κB-motifs underlined), was generated in an overlap-PCR using the LTR of Indie_C1 as the template. The additional 22 bp sequence constituting the F-κB motif was adopted from the HIV-1C molecular clone BL42-02 (GenBank accession No. HQ202921). The amplified FHHC-LTR was inserted into pcLGIT vector, substituting the original 3’-LTR. Subsequently, using the overlap-PCR, inactivating point mutations were introduced sequentially into the ‘FHHC’ (4-κB) LTR, to generate the other members of the panel: OHHC (3-κB), OOHC (2-κB), OOOC (1- κB) and OOOO (0-κB) (Figure 2A). Of note, the inactivation mutations only introduced base substitutions, not deletions, keeping the length of the viral promoter constant among the variant viral vectors. The mutated κB-motif ‘O’ contains the sequence <u>TCT</u>ACTTT<u>TT</u> (underlined bases represent inactivating mutations). The variant LTR fragments were cloned directionally between the XhoI and PmeI sites present on the outer primers-N1990 FP (5’- GCGTACCTCGAGTGGAAGGGTTAATTTACTCCAAGAAAAGGC-3’) and N1991 RP (5’-TATGTCGTTTAAACCTGCTAGAGATTTTCCACACTACCAAAAGGGTCTGAG-3’) thus, substituting the original 3’-LTR of pcLGIT. Therefore, the members of the vector panel are genetically identical except for the differences in the functional NF-κB motifs in the LTR. The internal primer sequences to generate the point mutations in the LTR variants are mentioned in Table 1. The 3’ LTR sequences of all the panel members were sequence-confirmed, and the expression of EGFP was ascertained in HEK293T cells.

**Table 1:**
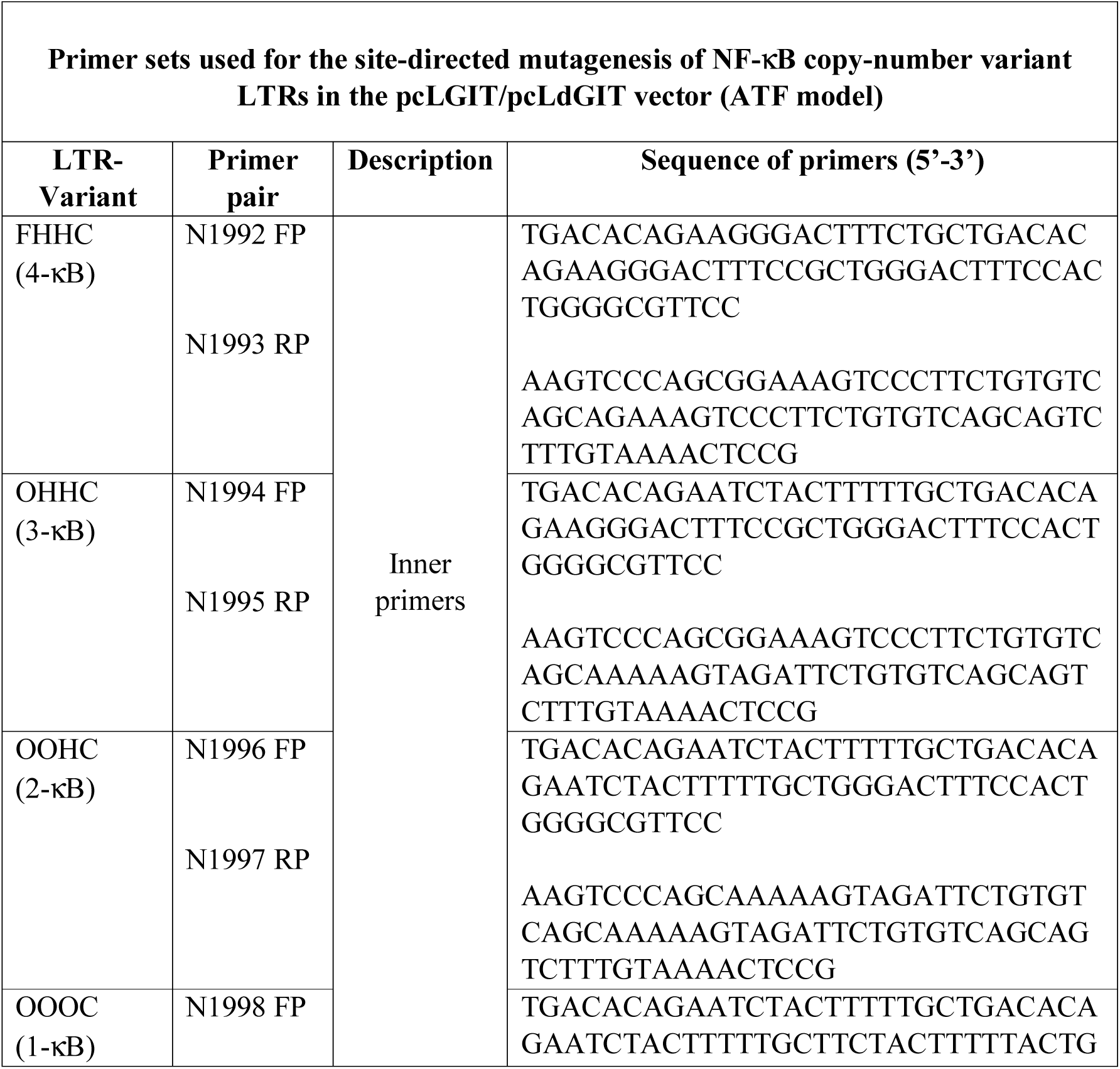
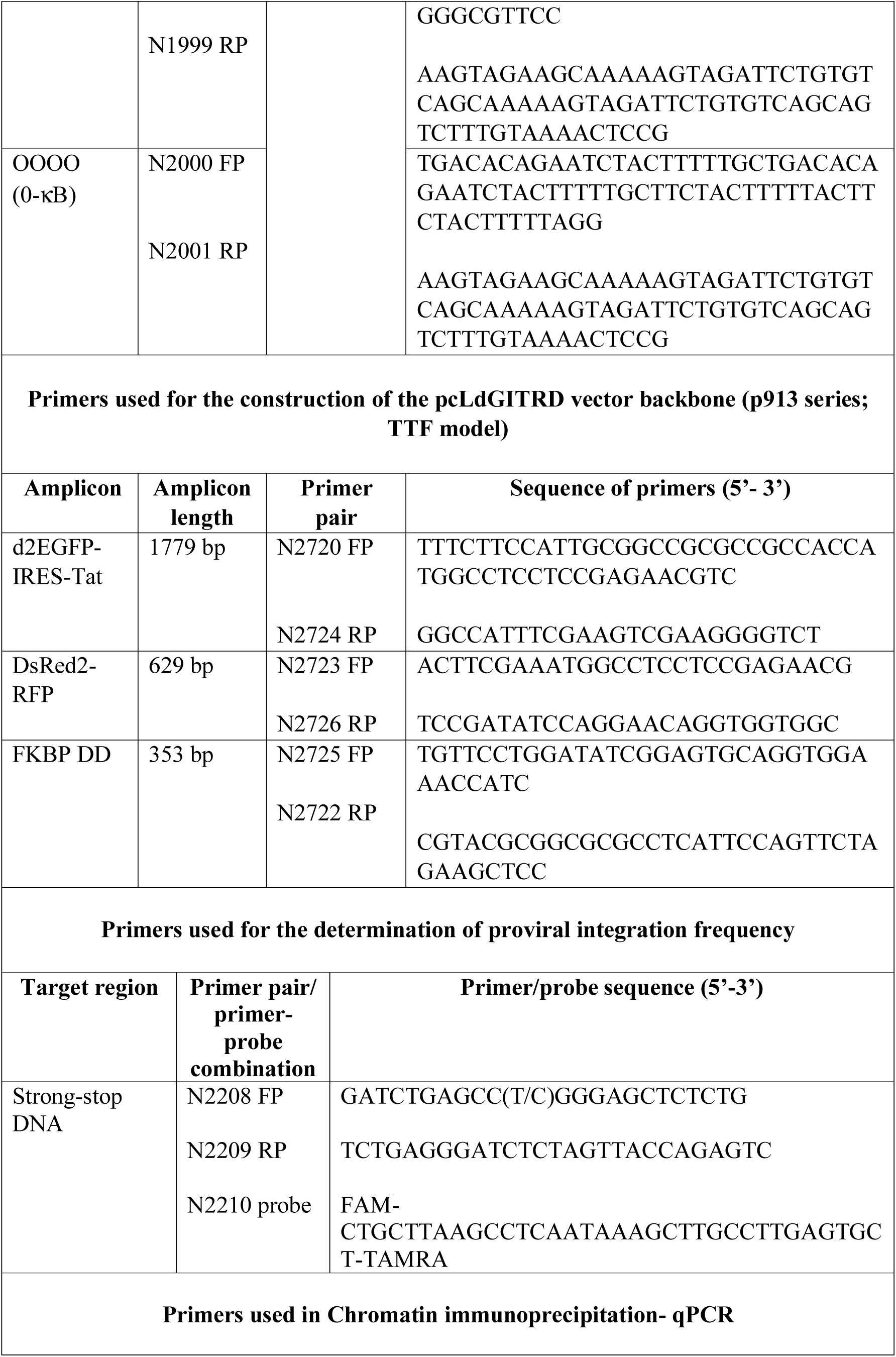
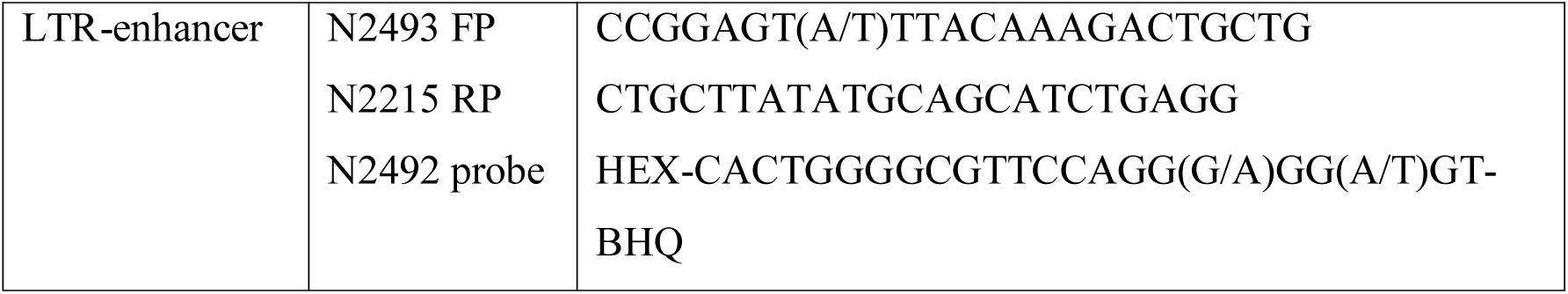
Primer sets used in PCR

A second panel of the five variant viral vectors, analogous to the p911a panel was also constructed using the pcLdGIT backbone, where EGFP was substituted with d2EGFP, a variant form of the fluorescent protein characterized by the shorter half-life (p911b vector series). First, the d2EGFP ORF was amplified from the pCAG-GFPd2 plasmid (#14760, Addgene, Massachusetts, USA) using the primer pair- N1142 FP (5’- CAGGAATTCGATGCTACCGGTCGCCACCATG-3’) and N1143 RP (5’- TCCTACTAGTAGGATCTGAGTCCGGACTACACATTGATCC-3’) and directionally cloned between the AgeI and BspE1 sites to replace the EGFP with the d2EGFP reporter gene in the pcLGIT backbone. To generate the p911b panel, the variant LTRs of the p911a panel were transferred directionally to the pcLdGIT vector backbone between the PmeI and XhoI sites, thus substituting the original 3’-LTR. The expression of d2EGFP from all the vectors of the p911b panel was verified using HEK293T cells.

#### Tunable Tat-feedback (TTF) model

In the TTF model, HIV-1C LTR regulates the co-expression of d2EGFP and Tat-RFP fusion protein from the vector pcLdGITRD (cLTR-d2EGFP-IRES-cTat:RFP:DD). The 5’ LTR in the pcLdGITRD vector transcribes a single transcript encoding d2EGFP and a 1,314 bp long fusion cassette separated by an IRES element. The fusion cassette is a combination of three different ORFs- (i) the cTat expression segment (BL4-3, GenBank accession number FJ765005.1), (ii) the ORF of DsRed2-RFP, and (iii) the FKBP destabilization domain (DD) (28). The three components of the ‘Tat:RFP:DD’ cassette were independently amplified using appropriate templates and primers, and, finally, using an overlap PCR, the fusion ORF was generated (primer sequences provided in Table 1). The Tat ORF from the pcLdGIT-3-κB vector (p911b series; ATF model) was replaced with the ‘Tat:RFP:DD’ ORF, thus, generating the pcLdGITRD-3-κB viral vector. pcLdGITRD-3-κB was subsequently used as the parental vector to construct the other member-pcLdGITRD-1-κB of the panel p913 (Figure 6A) by cloning the respective 3’LTRs between PmeI and XhoI in the pcLdGITRD backbone. The d2EGFP expression from the two members of the panel p913 was confirmed in HEK293T cells.

### Shield1 dose-dependent Tat:RFP:DD expression from the pcLdGITRD vector (TTF model) in HEK293T cells

The FKBP DD-tag in the pcLdGITRD construct marks the ‘Tat:RFP:DD’ fusion protein for rapid degradation through the proteasome pathway (28). However, ‘Shield1’, a 750 kD cell-permeable ligand can bind the DD motif and rescue the fusion cassette from rapid processing in a dose-responsive manner with minimum off-target effects (29). Thus, by changing the concentration of Shield1 in the culture medium, the pcLdGITRD construct permits fine-tuning of the intracellular concentration of Tat without altering the transcriptional strength of the LTR. To validate the Shield1 dose-dependent Tat:RFP:DD expression, approximately 0.6 million HEK239T cells were transfected with 1 μg pcLdGITRD-3-κB vector in each well of a 12-well culture dish and treated with varying concentrations (0, 0.5, 1.0, 2.5, 4.0 and to 5.0 μM) of Shield1 (#632189, Takara Clontech). After 48 h of transfection, the expressions of both DsRed2-RFP and d2EGFP were recorded using a fluorescent microscope.

### Generation of pseudotyped reporter virus and the estimation of relative infectious units (RIU)

Pseudotyped reporter viruses were generated in HEK293T cells. Each viral vector was transfected together with the 3^rd^ generation lentiviral packaging vectors using the standard calcium phosphate protocol (82). Briefly, a plasmid DNA cocktail consisting of 10 μg of individual viral vector (NF-κB motif variants), 5 μg psPAX2 (#11348; NIH AIDS reagent program, Maryland, USA), 3.5 μg pHEF-VSVG (#4693; NIH AIDS Reagent program) and 1.5 μg pCMV-rev (#1443; NIH AIDS Reagent program) was transfected in a 100 mm dish seeded with HEK293T at 30% cell confluence. pCMV-RFP (0.2 μg) was used as an internal control for transfection. Six hours post-transfection, the medium was replenished with complete DMEM. Culture supernatants were harvested at 48 h post-transfection, filtered using 0.22 μ filter and stored in 1 ml aliquots in a deep freezer for future use.

The RIU of the pseudotyped reporter viruses was quantified in Jurkat T-cells by measuring EGFP or d2EGFP expression by flow cytometry. Precisely, 3 x 10^4^ Jurkat cells in each well of a 12-well tissue culture plate were infected with individual viral stocks serially diluted 2- fold (from 10 xd to 80 xd) in a total volume of 1 ml of 10% RPMI containing 25 μg/ml of DEAE-Dextran. Six hours post-infection, the cells were washed and replenished with 1 ml of complete RPMI. Post 48 h, the cells were activated with a combination of 40 ng/ml PMA (P8139, Sigma Aldrich), 40 ng/ml TNFα (T0157, Sigma-Aldrich), 200 nM TSA (T8552, Sigma Aldrich) and 2.5 mM HMBA (224235, Sigma-Aldrich) for 18 h, following which the percent EGFP^+^ or d2EGFP^+^ cells were analyzed using a flow cytometer (BD FACSAria III sorter, BD biosciences, New Jersey, USA). Following this, titration curves were constructed and analyzed for 5-10% infectivity of the cells by regression analysis, which would correspond to ∼0.05-0.1 RIU. For the TTF model, cells were maintained in 1μM Shield1 throughout the procedure.

### Viral gene-expression analysis

Viral gene-expression levels were compared among the LTR variant strains of the pcLGIT panel (ATF model). Towards this, approximately one million Jurkat cells were infected with each viral strain of the panel independently at an RIU of ∼0.5–0.6. Three days following infection, the cells were activated using a cocktail of global T-cell activators (40 ng/ml PMA + 40 ng/ml TNFα + 200 nM TSA + 2.5 mM HMBA) and 24 hours following activation, EGFP fluorescence (mean fluorescence intensity or MFI) was estimated for both the uninduced and activated cells using flow cytometry. Tat transcript levels of the control and activated cells, were determined using a Tat RT-PCR (see below). Fold enhancements in EGFP, and Tat expression levels were obtained from the ratios of the EGFP-MFI values of the activated and control samples and similarly their relative Tat-mRNA values, respectively.

To compare the gene-expression levels from the LTR at different transcriptional strengths of Tat-feedback, modulated by the Shield1 concentrations (TTF model), approximately 0.3 million Jurkat cells in a 35 mm culture dish were infected with the cLdGITRD-3-κB viral strain at a p24 equivalent of 20 ng/ml. Twenty-four hours post-infection, the infected cells were washed and replenished with complete RPMI. The cells were then equally distributed into four wells of a 6-well plate and each well treated with either 0, 0.5, 2.5, or 5.0 μM Shield1. After 48 hours, half of the cells from each Shield1 treatment were activated using the global T-cell activators, and 24 hours later, both the uninduced and activated fractions from each dose of Shield1 were subjected to d2EGFP and Tat-transcript expression analysis using flow cytometry and RT-PCR, respectively.

### FACS sorting and the generation of stable Jurkat cells and clonal lines

#### ATF model

Individual, NF-κB variant, pseudotyped viral stocks of the cLGIT/cLdGIT panel were added to 1 x 10^6^ Jurkat cells in a 35 mm culture dish, at an RIU of ∼0.05-0.1 in a total volume of 2 ml complete RPMI supplemented with 25 μg/ml DEAE-Dextran. After six hours of infection, cells were washed to remove DEAE-Dextran and transferred to 5 ml of complete RPMI medium in a T-25 flask and maintained under standard culture conditions.

The infected cell pools were expanded over seven days and induced with the global T-cell activation cocktail as mentioned above. After 18 h of activation, total EGFP^+^ (MFI >10^3^ RFU), EGFP^High^ (MFI >10^4^ RFU), or d2EGFP^High^ (MFI ∼0.5 × 10^3^ – 0.5 × 10^4^) cells were FACS sorted from respective cell pools depending on the subsequent viral latency assays. A small aliquot of the sorted cell population was re-analysed to confirm the purity of the sorted cells. Each sorted cell pool with a stable EGFP/d2EGFP expression represented a mixed population with random proviral integrations with the corresponding NF-κB variant strain.

Stable clonal cell lines of the cLGIT variant panel (expressing EGFP) were established by sorting a single cell per three wells in a 96-well plate. Each well of the collection plate contained 100 μl of a mix of equal proportions of complete- and the spent-RPMI media. The cells were diluted to a cell density of 0.1 x 10^6^/ml before the sort.

#### TTF model

1 x 10^6^ Jurkat cells were infected with the sub-genomic cLdGITRD viral strains (3- and 1-κB variants) at an RIU of ∼0.1-0.2 in 1 ml of complete RPMI medium supplemented with 25 μg/ml of DEAE-dextran and 1.0 μM Shield1. Six hours post-infection, the infected cells (J-cLdGITRD) were washed and replenished with 1 ml of complete RPMI supplemented with 1 μ Shield1. Next, 72 h following the infection, the cells were induced with the previously mentioned T-cell activation cocktail for 18 h, and following the activation, the stable, d2EGFP^+^ J-cLdGITRD cells were sorted.

### The analysis of proviral integration frequency

A Taqman qPCR was used to determine the mean number of proviral integrations per cell using the genomic DNA extracted from the stable, J-cLGIT cell pools, and also from the EGFP^High^ and EGFP^-^ subfractions corresponding to the representative bimodal clones of the 3-κB and the 4-κB variants. Genomic DNA was extracted from 1 x 10^6^ stable cells using the GenElute mammalian genomic DNA kit (G1N350, Sigma-Aldrich) following the manufacturer’s instructions. The extracted DNA was dissolved in TE, and the concentration was adjusted to 70 ng/μl. Five μl of this solution was equivalent to approximately 10^5^ copies of the human genome. The stock DNA solution was subjected to a 10-fold serial dilution up to a final DNA concentration of 10^1^ copies/5 μl and used as the template in the PCR. A 129 bp fragment spanning the R-U5 region of the HIV-1 5’ LTR (+18 to +147) was amplified using the primer-probe combination N2208 FP, N2209 RP, and N2210 FAM (see Table 1 for the primer sequences) in a Taqman real-time PCR. A standard curve was established simultaneously using the genomic DNA extracted from the J-Lat 8.4 cells that contain a single proviral copy per cell (78). The proviral copy number of the query samples was then estimated using the regression analysis.

### Generation of kinetic profiles of latency-establishment

#### ATF model

Kinetic curves of latency were established to compare the rates of promoter silencing among the NF-κB variant strains of the ATF panel. Towards this, the sorted cell pools (total EGFP^+^, EGFP^High,^ or d2EGFP^High^) were maintained under standard experimental conditions while a small aliquot was collected at regular intervals to monitor the EGFP/d2EGFP expression using the FACSAria III flow cytometer. Temporal kinetic profiles for % GFP^+^ cells and GFP-MFI were constructed and compared among the five NF-κB variants for the total EGFP^+ve^, EGFP^High^ as well as the d2EGFP^High^ cells. The EGFP reporter gene having a longer half-life (∼48 h) was measured every 4 days for 16-24 days while the analysis of d2EGFP expression with a shorter half-life (∼ 2h) was performed every 24 h following sorting, for 7 days.

Kinetic profiles of latency were established for the clonal lines of the cLGIT panel. The sorted, single EGFP^High^ cells corresponding to each NF-κB variant strain were expanded to form clonal lines, and 16 such lines corresponding to each cLGIT viral variant were flow-analyzed for their EGFP expression profiles on Days 21 and 28 post sorting.

#### TTF model

Latency-establishment profiles at varied strengths of the Tat-feedback circuit were studied using the TTF model by altering the Shield1 concentrations in the culture medium. The sorted d2EGFP^High^ cells corresponding to the cLdGITRD-3-κB (strong) and the cLdGITRD-1-κB (weak) LTR variants were divided into four separate fractions and maintained at four different concentrations of Shield1 (0, 0.5, 1 and 3 μM) under standard experimental conditions, while a small aliquot from all the fractions was collected every 24 h to monitor the expression of both d2EGFP and Tat-RFP using FACSAria III flow cytometer, for 7 days. Temporal kinetic profiles were constructed and compared among the two NF-κB variants as well as across the different concentrations of Shield1.

### Live-dead analysis of cells

A live-dead, viability assay was performed to exclude the dead cells before every flow analysis. Cell samples were stained using the LIVE/DEAD^TM^ fixable Red Dead Cell Stain Kit (#L34972, Molecular Probes, Thermo Fisher Scientific, Massachusetts, USA) following the manufacturer’s protocol. Briefly, cell samples were harvested in a microcentrifuge tube, washed once with 1X PBS, and resuspended in 500 μl of 1:1,000 diluted live-dead stain. The cells were then incubated for 30 mins at room temperature in the dark, following which they were washed, resuspended in 500 μl of RPMI supplemented with 2% RPMI and analyzed in the flow cytometer.

### DNA cell-cycle analysis

Cell cycle analysis by quantitating DNA content was performed on the EGFP^High^ and EGFP^-^ subfractions of the bimodal clonal lines of the J-cLGIT-3- and 4-κB variants. The standard propidium iodide (PI) staining protocol was followed with slight modifications to minimize the quenching of the EGFP signal (83). Briefly, 1 × 10^6^ cells were harvested in 1.5 ml microcentrifuge tubes and washed once with 1X PBS. The pellets were resuspended thoroughly in 1 ml fix solution (2% glucose and 2% paraformaldehyde in 1X PBS) and incubated on ice for 10 mins. The cells were then washed once with 1X PBS, resuspended in 100 μl of 1X PBS, and 900 μl of ice-cold 70% ethanol added dropwise with gentle vortexing. The fixed cells were incubated on ice for 1 h, washed with 1X wash solution (20 mM Hepes, 0.25% NP-40, 0.1% BSA in 1X PBS), supernatants aspired and resuspended in 500 μl of 1X PBS containing 10 μg/ml RNAse A and 20 μg/ml PI. This was followed by incubation in the dark for 30 mins at room temperature and analysis by flow cytometry. Pulse processing (pulse area vs pulse height) was used to exclude doublets, and debris and the gated singlets were applied on the PI histogram plot to determine the % cells in the G1, S, and G2/M phases.

### Quality assurance and data analysis in flow cytometry

The BD FACSAria III sorter was optimized and calibrated before every operation (flow analysis or sorting) to ensure quality performance. The instrument was calibrated for optical laser alignment, fluorescence and light scatter resolution, and fluorescence detector sensitivity using the BD^TM^ CS&T beads (#656504, BD Biosciences). Drop delay before every FACS sort was determined using the BD FACS^TM^ Accudrop beads (#345249, BD Biosciences) while fluorescence linearity and doublet discrimination before DNA cell-cycle analysis were assessed using the BD^TM^ DNA QC particles (#349523, BD Biosciences). Further, uniform PMT voltage parameters were set for every fluorescent channel using appropriate negative control samples to avoid variations in the MFI measurements at different time-points of the latency kinetics experiments.

All the flow cytometry data analyses were performed using FCS Express 4 and 6 versions (De Novo Software, Los Angeles, CA).

### The analysis of the Tat-transcripts in stable Jurkat cells

We quantitated Tat transcript levels using a real-time PCR as a surrogate marker of the transcriptional status of the LTR during latency-establishment, or latency reversal. Total mRNA was extracted from 0.5 x 10^6^ cells using a single-step RNA isolation reagent- TRI reagent (T9424, Sigma-Aldrich) at specified time points. Using random hexamer primers, 250-1,000 ng of extracted RNA was converted to cDNA in a reaction volume of 20 μl using the Tetro cDNA synthesis kit (BIO-65043, Bioline, London, UK). The cDNA was then amplified using an SYBR green RT-PCR kit (06924204001, Roche Products, Mumbai, India) for a 139 bp region in the Tat exon-1 using the primers- N1783 (5’- GGAATCATCCAGGAAGTCAGCCCGAAAC-3’) and N1784 (5’- CTTCGTCGCTGTCTCCGCTTCTTCCTG-3’). The GAPDH RT-PCR was employed as an internal control (primer pair N2232: 5’-GAGCTGAACGGGAAGCTCACTG-3’ and N2233: 5’- GCTTCACCACCTTCTTGATGTCA-3’). The relative gene expression was calculated using the ΔΔCt method.

### Indirect immunofluorescence of Tat

Immunofluorescence staining of Tat was performed at multiple time points during the establishment of viral latency in stable J-cLdGIT-3-κB cells characterized by strong d2EGFP fluorescence (MFI range 5 x 10^3^ to 50 x 10^3^). The sorted d2EGFP^High^ cells were considered as the D0 sample, and Tat-IF was performed subsequently at an interval of every 4 days. Approximately 3 x 10^6^ cells were collected in a 1.5 ml vial, washed once with 1X PBS, and fixed with 2% paraformaldehyde in PBS for 10 min at room temperature with mild rocking. Fixed cells were re-washed with 1X PBS followed by permeabilization with 0.2% Triton-X-100 in PBS for 10 min with gentle and intermittent vortexing. Fixed and permeabilized cells were then washed again with 1X PBS and blocked with 4% BSA in PBS for 30 min at room temperature with mild rocking. The blocked cells were incubated with a rabbit, polyclonal anti-Tat antibody (ab43014, Abcam, Cambridge, UK) at 1: 250 dilution for 1h at room temperature followed by two PBS washes. This was followed by the incubation with 1: 500 dilution of Goat anti-rabbit Alexa Fluor 568 (A-11010, Molecular Probes) for 20 min in the dark at room temperature followed by a PBS wash. The nucleus was stained with 4 μg/ml of DAPI for 20 min in the dark at room temperature. Cells were washed twice and mounted on coverslips with 70% glycerol for confocal imaging. Images were acquired with a Zeiss LSM 880 confocal laser scanning microscope with Airyscan using a Plan Apochromat X63/1.4- oil immersion objective and analyzed using the ZEN 2.1 software. For imaging single cells, a 4X higher zoom was applied. The fluorescent cut-off for d2EGFP expression was determined using uninfected Jurkat control, while that for Tat-Alexa 568 expression was set using the infected but no-primary Tat antibody control. At each time-point, 150 cells were analyzed; fluorescent intensities (AU) for d2EGFP and Tat-Alexa 568 expressions independently determined for the nuclear and extra-nuclear compartments of the cells and temporal curves generated.

### The proximity ligation assay

We used an in situ proximity ligation assay (PLA) to detect Tat in HEK293 cells independently infected with the cLdGIT-3- and 4-κB reporter virus. The assay was performed using a commercial kit (Duolink In Situ Red Starter kit Mouse/Rabbit, #DUO92101, Sigma-Aldrich) following the instructions of the manufacturer. Briefly, a heterogeneous population of HEK293 cells harboring both active and latent virus (cLdGIT-3- κB or 4-κB), marked by the presence or absence of green fluorescence, respectively, were seeded on glass coverslips and allowed to grow to 60-70% confluence. The evenly distributed cells on the coverslip were fixed with 4% paraformaldehyde for 20 min at room temperature, permeabilized with 0.1% Triton-X-100 for 10 min at room temperature and washed thrice with 1X PBS. This was followed by blocking for one hour using the reagent supplied in the kit. The blocked cells were then treated with the rabbit polyclonal anti-Tat antibody at 1: 250 dilution (Catalog no. ab43014, Abcam) in combination with the mouse monoclonal anti-Tat antibody at 1: 250 dilution (Catalog no. 7377, NIH AIDS reagent program, Maryland, USA). The cells were incubated with a pair of probes (the PLA probe Anti-Mouse MINUS; DUO92004 and PLA probe Anti-Rabbit PLUS; DUO92002) in a 40 μl reaction volume, for one hour at 37°C followed by washing twice with 500 μl of wash buffer A for 5 min each time. The ligation and amplification reactions were performed as per manufacturer’s instructions using the Duolink In Situ Detection reagents Red (Catalog no. DUO92008). The DAPI-supplemented mounting medium (Catalog no. DUO82040, supplied in the PLA kit) was used for mounting the cells. No-primary and single antibody controls were used to assess non-specific PLA spots. Imaging of the cells was performed using a Zeiss LSM 880 confocal laser scanning microscope with Airyscan fitted with a Plan Apochromat 63X/1.4 oil immersion objective. Signal intensities of the PLA positive spots were quantitated manually using the Image J software. A total of 164 cells were analyzed from the d2EGFP^+^ category (128 from the 4-κB and 36 from the 3-κB sample) while 168 cells were analyzed from the d2EGFP^-^ category (123 from the 4-κB and 45 from the 3-κB sample) to compare the number of PLA dots per cell between the two phenotypes.

The primary antibody pair, the rabbit polyclonal anti-Tat (ab43014, Abcam) and mouse monoclonal anti-Tat (7377, AIDS reagents program) antibodies, was validated for Tat specificity before performing PLA in stable HEK293 cells harboring the cLdGIT-3- and 4-κB proviruses. Approximately 0.5 x 10^6^ HEK293T cells in each well of an 8- micro chambered glass slide (80826, ibidi, Grafelfing, Germany) were transfected with either 800 ng of pcLGIT vector (B-Tat) or 200, 400 or 800 ng of pcLGIT vector (C-Tat). After 48 hours of transfection, Tat-PLA was performed as detailed above. Confocal images were captured using the same model of a confocal microscope and identical parameters, as mentioned above. Image J software was used to measure d2EGFP intensity (AU) and manual quantitation of Tat-PLA spots from 25 cells corresponding to each dose of pcLGIT vector.

### Chromatin immunoprecipitation assay

We used a chromatin preparation equivalent of 2 x 10^6^ cells (either EGFP^High^ or EGFP^-ve^) for each immunoprecipitation assay, as described previously (32). Briefly, 2 x 10^6^ Jurkat cells collected in a 1.5 ml vial were washed with 1X PBS, resuspended in 1 ml of RPMI supplemented with 1% formaldehyde and incubated with gentle agitation for 10 min at room temperature. The cross-linking reaction was quenched by incubating the cells with 0.125 M glycine for 5 min with mild agitation at room temperature followed by centrifugation at 3,000 rpm for 5 min at 4°C with a subsequent PBS wash (containing 0.01X protease inhibitor cocktail or PIC; #11836170001, Roche Applied Science, *Indianapolis*, USA). Following the complete removal of PBS, the cells were resuspended in 100 μl of ice-chilled lysis buffer (1% SDS, 50 mM Tris buffer, pH 8.0, 10 mM EDTA) and incubated on ice for 20 min with occasional mixing of the lysate using a wide-bore tip. The lysate in each vial was subjected to 22 cycles of sonication at the high mode, using 30-second-ON followed by a 30-second-OFF pulse scheme in the Bioruptor plus sonicator (UCD-300, Diagenode, Liege, Belgium) containing pre-chilled water. The sonicated lysate was centrifuged at 12,000 rpm for 10 min at 4°C to remove any cellular debris; the clear supernatant was transferred to a fresh 1.5 ml vial and stored at −80°C until use. One-tenth of the lysate (10 μl) was used to confirm the shearing of chromatin to generate 200-500 bp fragment sizes. Each IP comprised of 100 μl of lysate and 2 μg of an antigen-specific antibody against p50 (ab7971, Abcam, Cambridge, UK), p65 (ab7970, Abcam), NFAT1 (ab2722, Abcam), NFAT2 (ab2796, Abcam), HIV-1 Tat (ab43014, Abcam or #7377, NIH AIDS reagent program or #4374, NIH AIDS reagent program), RNA Pol II CTD phospho S2 (ab5095, Abcam), or H3K9 Tri Meth (ab8898, Abcam). The ChIPed DNA was amplified using the primer pair N1054 FP (5’- GATCTGAGCC(T/C)GGGAGCTCTCTG-3’) and N1056 RP (5’- TCTGAGGGATCTCTAGTTACCAGAGTC-3’) spanning a 240 bp sequence within the enhancer-core promoter region in the LTR. The amplified DNA fragments were subjected to agarose gel electrophoresis, and the band intensities were normalized using the percent-input method to compare differential recruitment of each transcription factor at the active vs. latent promoter. To enhance the sensitivity of the assay, TaqMan qPCR was performed using the ChIP-DNA and the primer-probe combination- N2493 FP, N2215 RP, and N2492 Hex (refer to Table 1). The final data were evaluated using the percent input method.

### Statistics

Statistical analyses were performed using GraphPad Prism 5.0 software. The statistical tests used to calculate *P* values are indicated in the corresponding figure legends.

## Acknowledgements

We thank Prof. Tapas Kumar Kundu (JNCASR, India) and Dr. Ravi Manjithaya (JNCASR, India) for intellectual discussions. We thank Dr. Uttara Chakraborty, S.L. Swaroopa Yalla and Dr. Narendra Nala of the flow cell at JNCASR, Suma B.S of the Confocal Imaging Facility and Anitha G. of the Sequencing Facility at JNCASR, India. We thank Neelakshi Varma and Surabhi Jirapure for initial help in establishing the latency models. Several reagents were obtained through the AIDS Research and Reference Reagent Program. This work was supported by grants to U.R. from the Department of Biotechnology (DBT), Government of India (DBT grant no. BT/PR7359/29/651/2012); National Institute of Health (NIH), USA (Grant No. NIDA 5RO1DA041751-02), and intramural funds from JNCASR.

## Author contributions

**S.C**., conception and design, acquisition of data, analysis and interpretation of data, drafting or revising the article, and providing essential unpublished data; **M.K.**, acquisition of data, analysis and interpretation of data, and providing essential unpublished data; **U.R.**, conception and design, fund acquisition, validation, writing, reviewing and editing the article.

